# A computational model of loss of dopaminergic cells in Parkinson’s disease due to glutamate-induced excitotoxicity

**DOI:** 10.1101/385138

**Authors:** Vignayanandam R. Muddapu, Alekhya Mandali, Srinivasa V. Chakravarthy, Srikanth Ramaswamy

## Abstract

Parkinson’s disease (PD) is a neurodegenerative disease associated with progressive and inexorable loss of dopaminergic cells in Substantia Nigra pars compacta (SNc). A full understanding of the underlying pathogenesis of this cell loss is unavailable, though a number of mechanisms have been indicated in the literature. A couple of these mechanisms, however, show potential for the development of radical and promising PD therapeutics. One of these mechanisms is the peculiar metabolic vulnerability of SNc cells by virtue of their excessive energy demands; the other is the excitotoxicity caused by excessive glutamate release onto SNc by an overactive Subthalamic Nucleus (STN). To investigate the latter hypothesis computationally, we developed a spiking neuron network model of the SNc-STN-GPe system. In the model, prolonged stimulation of SNc cells by an overactive STN leads to an increase in a ‘stress’ variable; when the stress in a SNc neuron exceeds a stress threshold the neuron dies. The model shows that the interaction between SNc and STN involves a positive feedback due to which, an initial loss of SNc cells that crosses a threshold causes a runaway effect that leads to an inexorable loss of SNc cells, strongly resembling the process of neurodegeneration. The model further suggests a link between the two aforementioned PD mechanisms: metabolic vulnerability and glutamate excitotoxicity. Our simulation results show that the excitotoxic cause of SNc cell loss in PD might be initiated by weak excitotoxicity mediated by energy deficit, followed by strong excitotoxicity, mediated by a disinhibited STN. A variety of conventional therapies are simulated in the model to test their efficacy in slowing down or arresting SNc cell loss. Among the current therapeutics, glutamate inhibition, dopamine restoration, subthalamotomy and deep brain stimulation showed superior neuroprotective effects in the proposed model.

## I. Introduction

There is a long tradition of investigation into the etiology and pathogenesis of Parkinson’s disease (PD) that seeks to link molecular (pesticides, oxidative stress, protein dysfunction etc) and subcellular (mitochondrial dysfunction etc) factors with the disease development. However, recent years see the emergence of two novel lines of investigation into PD pathogenesis. These approaches, that aim to understand the PD pathology at cellular and network level, mark a significant deviation from the traditional approaches (Rodriguez et al., 1998; Bolam and Pissadaki, 2012; Pissadaki and Bolam, 2013; Balasubramani et al., 2015a, 2015b; Pacelli et al., 2015; Mandali and Chakravarthy, 2016; Chakravarthy and Moustafa, 2018).

One of these approaches believes that the primary factors that cause degeneration of dopaminergic cells of SNc are metabolic in nature. Since the metabolic demands of SNc neurons are particularly high, any sustained insufficiency in the supply of energy can result in cellular degeneration characteristic of PD (Mergenthaler et al., 2013). According to the second approach, the overactivity of STN in PD causes excessive release of glutamate to SNc, which in turns causes degeneration of SNc neurons by glutamate excitotoxicity (Rodriguez et al., 1998). Even the above two approaches are interrelated and not totally independent since one form of excitotoxicity – the ‘weak excitotoxicity’ – is thought to have its roots in impaired cellular metabolism (Albin and Greenamyre, 1992).

Therefore, at the bottom of these new lines of investigation of PD pathogenesis, is the insight that the mismatch in energy supply and demand could be a primary factor underlying neurodegeneration in PD. Such a mismatch is more likely to take place in special nuclei like SNc due to their peculiar metabolic vulnerability. Similar ideas have been proffered to account for other forms of neurodegeneration like, for example, Huntington’s disease, Alzheimer’s disease, and amyotrophic lateral sclerosis also (Beal et al., 1993; Johri and Beal, 2012; Gao et al., 2017).

If metabolic factors are indeed the deep underlying reasons behind PD pathogenesis, it is a hypothesis that deserves a closer attention and merits a substantial investment of time and effort for an in-depth study. This is because any positive proof regarding the role of metabolic factors puts a completely new spin on PD research. Unlike previous therapeutic approaches that basically manage the disease, and hold no promise of a cure, the new approach can in principle point to a more lasting solution. If inefficient energy delivery or energy transformation mechanisms are the reason behind degenerative cell death in PD, relieving the metabolic load on the vulnerable neuronal clusters, by intervening through brain stimulation and/or pharmacology could prove to be effective treatments for PD, a disease that hitherto proved itself to intractable to these standard approaches.

### Parkinson’s Disease

Parkinson’s disease (PD) is the second most prominent neurodegenerative disease around the world next to Alzheimer’s disease. It is caused by loss of dopaminergic neurons in substantia nigra (German et al., 1989; Fearnley and Lees, 1991; Hindle, 2010) of basal ganglia which causes cardinal symptoms such as bradykinesia, tremor, rigidity, postural imbalance, freezing gait and other cognitive dysfunctions (Sveinbjornsdottir, 2016).

Basal ganglia (BG) play an important role in controlling motor movements through two distinct pathways – direct and indirect pathways. BG is situated in midbrain region which consists of seven nuclei namely substantia nigra (pars compacta (SNc) and pars recticulata (SNr)), globus pallidus (externa (GPe) and interna (GPi)), striatum (caudate and putamen), subthalamic nucleus (STN). The cortical inputs to the striatum (input nuclei of BG) flow in two ways - directly to GPi (output nuclei of BG) constitute the direct pathway and indirectly to GPi through GPe and STN constitute the indirect pathway. Activation of direct and indirect pathways results in initiation and inhibition of movements respectively. The balance between direct and indirect pathway is modulated by dopamine which is released by SNc (Chakravarthy and Moustafa, 2018).

To alleviate PD symptoms, Levodopa (L-DOPA) is given as dopamine supplement which is a precursor of dopamine. But the long-term treatment of L-DOPA results in side-effects such as L-DOPA-induced dyskinesias (LID). To overcome LID in advanced PD, deep brain stimulation (DBS) to STN is employed which reduces dyskinesias (Kim et al., 2015) and also increases the half-life of L-DOPA (Odekerken et al., 2013; Xie et al., 2016).

### Causes of SNc cell loss

The loss of SNc neurons in PD is not clearly elucidated in contemporary literature. It can be caused by genetic mutations, aging, inflammation and environmental toxins (Surmeier et al., 2010). Why are SNc neurons more vulnerable in PD compared to other neurons? It can be due to their complex axonal arborisation (Bolam and Pissadaki, 2012; Pacelli et al., 2015), presence of pacemaking ion channels (Dragicevic et al., 2015a), reactive neurotransmitter dopamine (Mosharov et al., 2009), glutamate excitotoxicity (Rodriguez et al., 1998), calcium loading and higher basal metabolic rate (Pacelli et al., 2015). There might be divergent causes as mentioned above, but all of them converge to common mechanisms such as oxidative stress, mitochondrial impairment and protein mishandling (Greenamyre and Hastings, 2004).

### Energy basis of neurodegeneration

In this paper, with the help of computational models, we investigate the hypothesis that the cellular energy deficiency in SNc is the major cause of SNc cell loss in PD. *The higher metabolic demand of SNc cells due to their unique molecular characteristics, complex morphologies, and other energy-demanding features perhaps make them more vulnerable to energy deficit*. Therefore, prolonged energy deprivation or insufficiency in such cells creates metabolic stress, eventually leading to neurodegeneration. If we can somehow reduce the metabolic stress on SNc cells, we can possibly delay the progression of cell loss in PD.

The hypothesis that systems-level energy imbalance is probably a key cause of PD was proposed by several researchers (Wellstead and Cloutier, 2011; Bolam and Pissadaki, 2012). According to (Wellstead and Cloutier, 2011), brain energy metabolism should be the core module for any modeling framework of PD to which other modules which explains cellular processes involved in PD pathogenesis can be incorporated to understand the etiology of PD. Bolam and Pissadaki (2012) emphasize the presence of complex axonal arborization that renders SNc neurons more susceptible in PD. They explore this idea using a computational model: as the size and complexity of axonal arborization increases, the metabolic cost of membrane potential recovery and propagation of action potential also increases (Pissadaki and Bolam, 2013). Due to their characteristic morphology, basal mitochondrial activity and oxidative stress were elevated, compared to normal cells as observed in ex *vivo* neuronal culture studies (Pacelli et al., 2015).

Albin and Greenamyre (Albin and Greenamyre, 1992) suggest that excitotoxicity caused by three mechanisms – strong excitotoxicity (due to excess excitatory neurotransmitter), receptor dysfunction (that causes increased susceptibility to excitatory neurotransmitter) and weak excitotoxicity (increased susceptibility to excitatory neurotransmitter due to impaired cellular energy metabolism). Here, we would like to explore “weak excitotoxicity hypothesis” where energy deficit eases magnesium blockage on NMDA receptors, results in the opening of ion channels and the subsequently causes neuronal damage.

In the proposed modeling study, we focus on excitotoxicity in SNc caused by STN which is precipitated by energy deficiency (Greene and Greenamyre, 1996) and exploring simulated therapeutic strategies for slowing down SNc cell loss. Since the mitochondrion is the source of energy for any cell, impairment in their mitochondrial function results in energy deficiency in the cell. It has been reported that 1-methyl 4-phenyl-1,2,3,6-tetrahydropyridine (MPTP) inhibits mitochondrial function (respiration) by inhibiting complex I (NADH ubiquinone oxidoreductase) (Tipton and Singer, 1993) which in turn results in the reduced reproduction of adenosine triphosphate (ATP). According to Rodriguez and co-workers, initial loss of dopamine in SNc leads to disinhibition and overactivity of the subthalamic nucleus (STN) which in turn causes excitotoxic damage to their target structures, including the SNc itself (Rodriguez et al., 1998). In addition, inhibition of mitochondrial activity by 1-methyl 4-phenyl-1,2,3,6-tetrahydropyridine (MPTP) or rotenone or 6-hydroxydopamine (6-OHDA) creates an energy deficit, which makes SNc neurons more susceptible to even physiological concentrations of glutamate release by STN (Albin and Greenamyre, 1992; Beal et al., 1993; Greene and Greenamyre, 1996; Rodriguez et al., 1998; Blandini, 2001, 2010; Ambrosi et al., 2014).

The pharmacological or surgical therapies that abolish the pathological oscillations in STN or block the receptors on SNc can be neuroprotective and might slow down the progression of SNc cell loss (Rodriguez et al., 1998). It has been reported that glutamate antagonists (Turski et al., 1991; Zuddas et al., 1992b) or STN ablation (Nakao et al., 1999) or long-term deep brain stimulation (DBS) of STN (Maesawa et al., 2004; Temel et al., 2006) results in survival of remaining SNc neurons in PD animal models.

With the help of computational models of neurovascular coupling, our group had earlier explored the effect of rhythms of energy delivery from the cerebrovascular system on neural function (Gandrakota et al., 2010; Chander and Chakravarthy, 2012; Chhabria and Chakravarthy, 2016; Philips et al., 2016). Recently, we proposed a preliminary computational spiking network model of STN-mediated excitotoxicity in SNc with a slightly abstract treatment of apoptosis (Muddapu and Chakravarthy, 2017). Simulation results showed that a sufficiently low initial value of a parameter known as the *stress threshold* (Saxena and Caroni, 2011) (analogous to the apoptotic threshold) causes initial SNc cell loss. This initial loss results in reduced dopamine content, which in turn leads to disinhibition of STN activity, eventually producing a runaway effect wherein SNc cells are lost due to excitotoxic damage. From model simulations, it was also shown that inhibition of STN activity may delay the progression of SNc cell loss. Building on the previous version of the excitotoxicity model, we have improved the excitotoxicity model and also explored the therapeutic strategies to slow down or halt the progression of SNc cell loss.

## II. Methods

All the nuclei were modeled as Izhikevich 2D neurons (*Figure-1*) All the simulations were performed by numerical integration using MATLAB 2016 with a time step (*dt*) of 0.1 sec.

**Figure 1:**
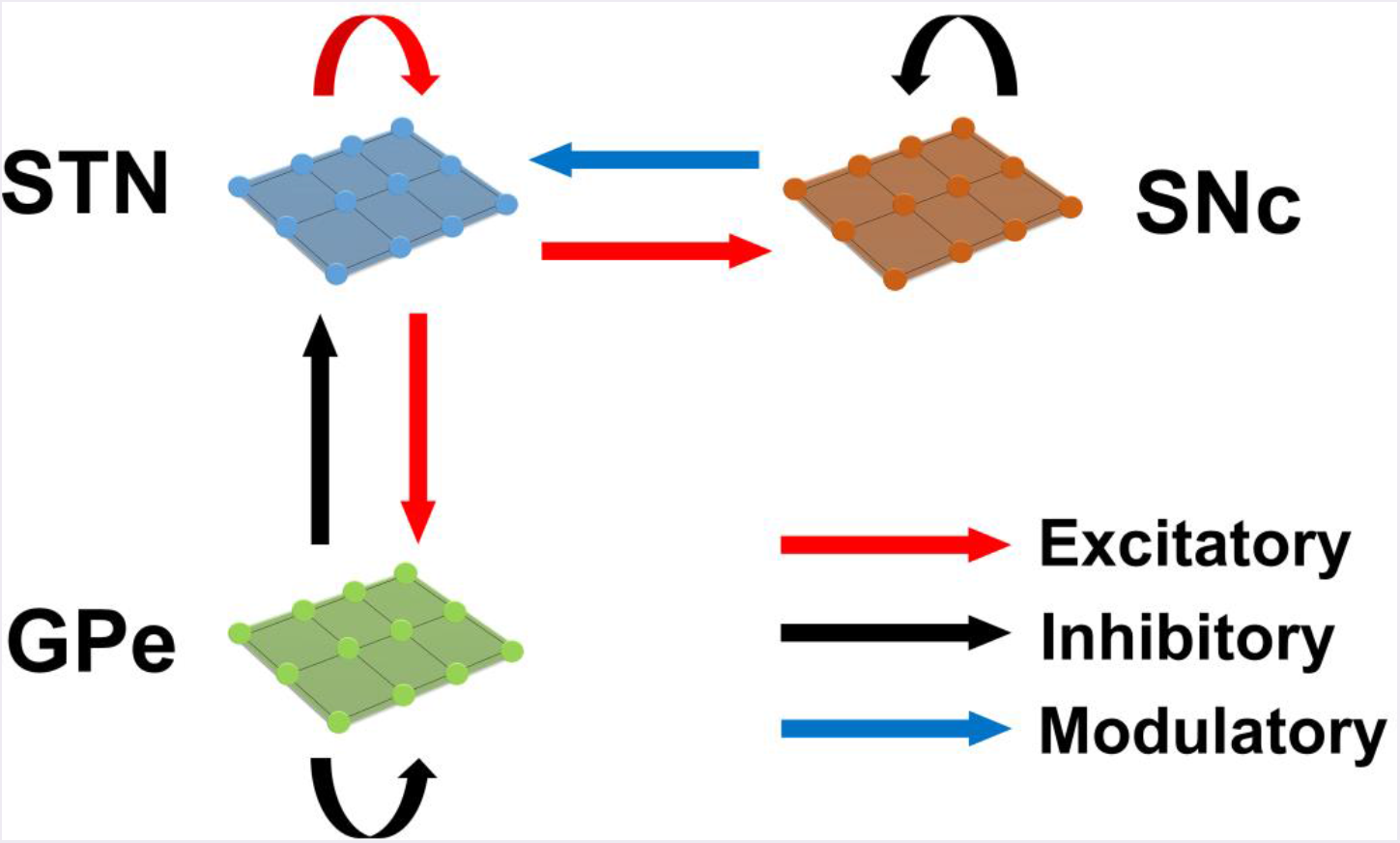
The model architecture of the proposed model of excitotoxicity.

### Izhikevich Neuron Model

Computational neuroscientists are often required to select the level at which a given model of interest must be cast i.e., biophysical-level, conductance-based modeling level, spiking neuron-level or rate-coded level. Biophysical models are biologically plausible but are computationally expensive whereas rate-coded models are computationally inexpensive but possess poor biological realism. To overcome this predicament, Izhikevich (Izhikevich, 2003) developed spiking neuron models that are computationally inexpensive and yet are able to capture various biological neuronal dynamics. The proposed model of excitotoxicity consists of SNc, STN, and GPe which were modeled using Izhikevich neuron models arranged in a 2D lattice (*Figure-1*) The population sizes of these nuclei in the model were selected based on the real numbers of neurons in these nuclei in rat basal ganglia (Oorschot, 1996). The Izhikevich parameters for STN and GPe were adapted from (Michmizos and Nikita, 2011; Mandali et al., 2015) and the parameters for SNc were adapted from (Cullen and Wong-Lin, 2015). The firing rates of these neuronal types were tuned to match the published data (Modolo et al., 2007; Tripathy et al., 2014) by varying the external bias current 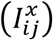. All parameters values are given in *Table 1*. The Izhikevich model consists of two variables, one for membrane potential (*v^x^*) and the other one for membrane recovery variable (*u^x^*)

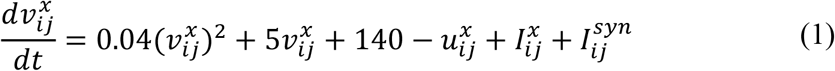

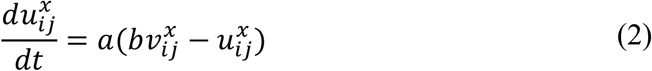

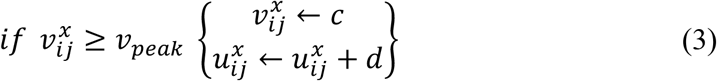

where, 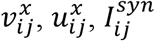 and 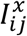 are the membrane potential, the membrane recovery variable, the total amount synaptic current received and the external current applied to neuron *x* at location (*i,j*) respectively, *v_peak_* is the maximum membrane voltage set to neuron (+30 mV) with *x* being GPe or SNc or STN neuron.

**Table-1.** Parameter values used in the proposed model

| Parameter(s) | STN | SNc | GPe |
| --- | --- | --- | --- |
| Izhikevich parameters ($a, b, c, d$) | $a = 0.005,$<br>$b = 0.265,$<br>$c = -65,$<br>$d = 1.5$ | $a = 0.0025,$<br>$b = 0.2,$<br>$c = -55,$<br>$d = 2$ | $a = 0.1,$<br>$b = 0.2,$<br>$c = -65,$<br>$d = 2$ |
| External current ($I^x$) | $I^{STN} = 3$ | $I^{SNc} = 9$ | $I^{GPe} = 4.25$ |
| Number of laterals ($nlat_x$) | $nlat_{STN} = 11$ | $nlat_{SNc} = 5$ | $nlat_{GPe} = 15$ |
| Radius of Gaussian laterals ($R_x$) | $R_{STN} = 1.4$ | $R_{SNc} = 1.6$ | $R_{GPe} = 1.6$ |
| Synaptic strength within laterals ($A_x$) | $A_{STN} = 1.3$ | $A_{SNc} = 0.1$ | $A_{GPe} = 0.1$ |
| Time decay constant for AMPA ($\tau_{AMPA}$) | 6 ms | 6 ms | 6 ms |
| Time decay constant for NMDA ($\tau_{NMDA}$) | 160 ms | 160 ms | 160 ms |
| Time decay constant for GABA ($\tau_{GABA}$) | 4 ms | 4 ms | 4 ms |
| Synaptic potential of AMPA receptor ($E_{AMPA}$) | 0 mV | 0 mV | 0 mV |
| Synaptic potential of NMDA receptor ($E_{NMDA}$) | 0 mV | 0 mV | 0 mV |
| Synaptic potential of GABA receptor ($E_{GABA}$) | -60 mV | -60 mV | -60 mV |
| Parameter that affects the post-synaptic current ($cd2$) | 0.1 | 0.1 | 0.1 |
| Concentration of Magnesium ($Mg^{2+}$) | 1 nM | 1 nM | 1 nM |

### Synaptic connections

The presence of excitatory synaptic connectivity from *STN* to *SNc* was observed from anatomical and electrophysiology studies (Hamani et al. 2004, 2017; Hammond et al. 1978; Iribe et al. 1999; Hitoshi Kita and Kitai 1987; Meissner et al. 2003; Mintz et al. 1986; Paquet et al. 1997; Smith and Grace 1992; Yoland Smith, Charara, and Parent 1996) and these connections might take part in controlling the bursting activity of SNc (Smith and Grace 1992). The number of neurons considered in SNc, STN and GPe populations are 64 (= 8 *x* 8), 1024 (= 32 *x* 32) and 1024 (= 32 *x* 32). A window of (4 *x* 4) STN neurons project to a single SNc neuron which was modeled as in (Humphries et al., 2009; Mandali et al., 2015). Similarly, the synaptic connectivity between *GPe* and *STN* was considered one-to-one as in (Dovzhenok and Rubchinsky, 2012; Mandali et al., 2015).

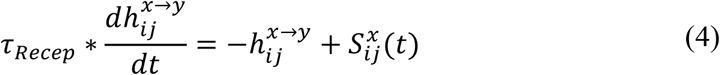

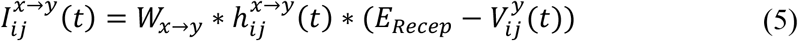

The *NMDA* currents are regulated by voltage-dependent magnesium channel (Jahr and Stevens, 1990) which was modeled as,

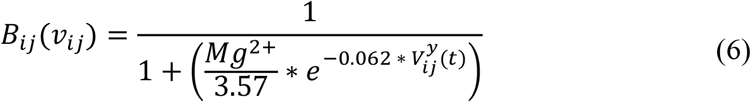

where, 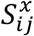 is the spiking activity of neuron *x* at time *t*, *τ_Recep_* is the decay constant for synaptic receptor, 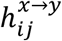 is the gating variable for the synaptic current from *x* to *y*, *W*_*x*→*y*_ is the synaptic weight from neuron *x* to *y*, *Mg*^2+^ is the magnesium ion concentration, 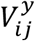 is the membrane potential of the neuron *y* for the neuron at the location (*i, j*) and *E_Recep_* is the receptor associated synaptic potential (*Recep* = NMDA/AMPA/GABA). The time constants of NMDA, AMPA and GABA in GPe, SNc and STN were chosen from (Götz et al., 1997) are given in *Table-1*.

### Lateral connections

The presence of lateral connections in STN (Kita, Chang, and Kitai 1983) and GPe (Kita and Kita 1994) were observed in various anatomical studies. The presence of GABAergic interneurons in SNc and their control of SNc activity was revealed by immunohistochemistry studies (Hebb and Robertson 1999; Tepper and Lee 2007). To simplify the model, GABAergic interneurons were replaced by GABAergic lateral connections in SNc population. Experimental studies show that synaptic current from lateral connections follows Gaussian distribution (Lukasiewicz and Werblin, 1990). The lateral connections in various modules in the current network (*STN, GPe and SNc*) were modeled as Gaussian neighborhoods (Mandali et al., 2015) and parameters used are given in *Table-1*. Each neuron receives synaptic input from a set number of neighboring neurons located in a 2D grid of size *nxn*.

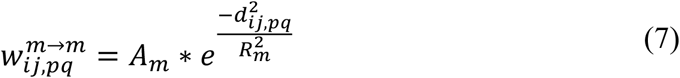

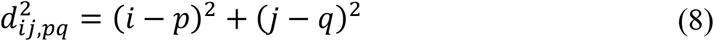

where, 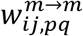 is the lateral connection weight of neuron type *m* at location (*i,j*), *d_ij,pq_* is the distance from center neuron (*p, q*), *R_m_* is the variance of Gaussian, *A_m_* is the strength of lateral synapse, *m* = *GPe or SNc or STN*.

### Effect of DA on synaptic plasticity

Several experimental studies demonstrate dopamine-dependent synaptic plasticity in STN (Hassani et al., 1997; Magill et al., 2001; Yang et al., 2016) and GPe (Magill et al., 2001; Mamad et al., 2015). Experimental observations show an increase in synchrony in STN (Bergman et al., 1994, 1998) and GPe populations (Bergman et al., 1998) at low DA levels. These conditions were implemented in the model by increasing in lateral connections strength in STN population as in (Hansel et al., 1995) and similarly decrease in lateral connections strength in GPe as in (Wang and Rinzel 1993) at low DA levels. Similarly, SNc populations also showed an increase in synchrony at low DA levels (Hebb and Robertson, 1999; Vandecasteele et al., 2005; Tepper and Lee, 2007; Ford, 2014) which was modeled similar to the model of DA-modulated GPe.

We modeled DA effect on the network as follows: as DA level increases, the strength of the lateral connections in STN decreases whereas, in GPe and SNc, lateral connections become stronger. These changes lead to a decrease in synchrony in all the 3 neuronal populations – GPe, SNc, and STN. Lateral strength was modulated by DA as follows,

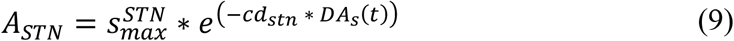

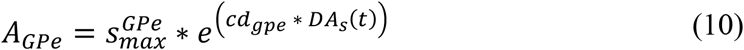

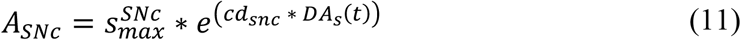

where, 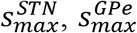 and 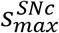 are strength of the lateral connections at the basal spontaneous activity of the population without any external influence in STN, GPe and SNc respectively. *cd_stn_*, *cd_gpe_* and *cd_snc_* were the factors by which dopamine affects the lateral connections in STN, GPe and SNc populations respectively.

According to experimental studies, DA causes post-synaptic effects on afferent currents in GPe and STN (Shen and Johnson, 2000; Smith and Kieval, 2000; Magill et al., 2001; Cragg et al., 2004; Fan et al., 2012). DA causes post-synaptic effects on afferent currents in SNc through somatodendritic DA receptors (Jang et al., 2011; Courtney et al., 2012; Ford, 2014). Thus, we included a factor (*cd2)*, which regulates the effect of DA on synaptic currents of GPe, SNc and STN. As observed in (Kreiss et al., 1997), as DA level increases, the regulated current decreases as follows:

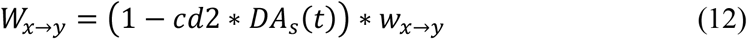

where, *W*_*x*→*y*_ is the synaptic weight (*STN* → *GPe*, *GPe* → *STN*, *STN* → *STN*, *GPe* → *GPe*,*STN* → *SNc*,*SNc* → *SNc*), *cd2* is the parameter that affects the post-synaptic current, *DA_s_*(*t*) is the instantaneous dopamine level which is the spatial average activity of all the neurons in SNc.

### Total synaptic current received by each neuron

#### STN

The total synaptic current received by a *STN* neuron at lattice position (*i,j*) is the summation of lateral glutamatergic input from other *STN* neurons considering both *NMDA* and *AMPA* currents and the GABAergic input from the *GPe* neurons.

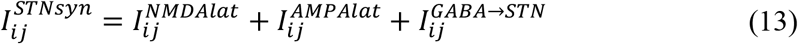

where, 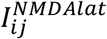 and 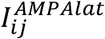 are the lateral glutamatergic current from other *STN* neurons considering both *NMDA* and *AMPA* receptors respectively, 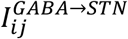 is the GABAergic current from *GPe* neuron.

#### GPe

The total synaptic current received by a *GPe* neuron at lattice position (*i,j*) is the summation of the lateral GABAergic current from other *GPe* neurons and the glutamatergic input from the *STN* neurons considering both *NMDA* and *AMPA* currents.

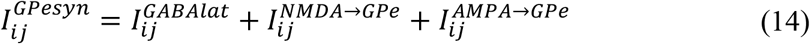

where, 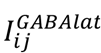 is the lateral GABAergic current from other *GPe* neurons, 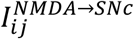 and 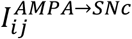 are the glutamatergic current from *STN* neuron considering both *NMDA* and *AMPA* receptors respectively.

#### SNc

The total synaptic current received by a *SNc* neuron at lattice position (*i,j*) is the summation of the lateral GABAergic current from other *SNc* neurons and the glutamatergic input from the *STN* neurons considering both *NMDA* and *AMPA* currents.

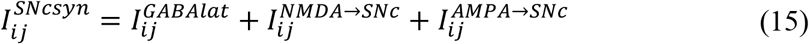

where, 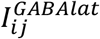 is the lateral GABAergic current from other *SNc* neurons, 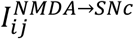 and 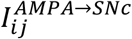 are the glutamatergic current from *STN* neuron considering both *NMDA* and *AMPA* receptors respectively.

### Neurodegeneration

According to (Rodriguez et al., 1998), dopamine deficiency in SNc leads to disinhibition and overactivity of the STN, which in turn causes excitotoxic damage to its target structures, including SNc itself. In order to simulate the SNc excitotoxicity induced by STN, we incorporate a mechanism of programmed cell death, whereby an SNc cell under high stress kills itself. The stress on a given SNc cell was calculated based on the firing history of the cell – higher firing activity causes higher stress.

The stress of each SNc neuron at lattice position (*i,j*) at time *t* due to excess firing is calculated as,

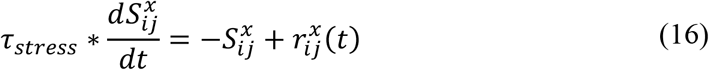

where, 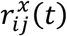 is instantaneous mean firing rate of a SNc neuron at lattice position (*i,j*) at time *t*, calculated with a fixed sliding window Δt (1 *sec*) (Grün and Rotter, 2010) as,

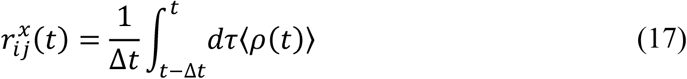

and

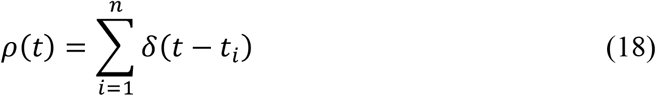

*Sequence of spike timing*: *t_i_* = 1,2,3 … ‥*n*

If stress variable 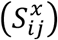 of a SNc neuron at lattice position (*i,j*) crosses certain threshold (*S_thres_*) then that particular SNc neuron will be eliminated (Iglesias and Villa, 2008).

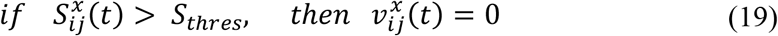

*Estimating rate of degeneration*

For a given course of SNc cell loss, the half-life is the time taken for half of the SNc cells to be lost (*t*_1/2_) The following equation was used to estimate the number of SNc cells *N_sc_*(*t*) for a given course that survived after a given time *t*.

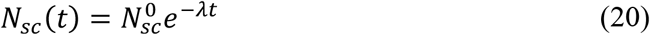

where, *λ* is the rate of degeneration (*sec*^1^), 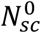 is the number of surviving SNc cells at *t* = 0

To estimate the rate of degeneration *λ* from a given course of SNc cell loss, the following equation was used,

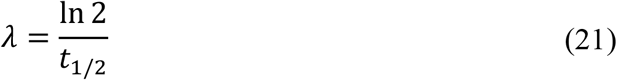

The instantaneous rate of degeneration *λ*(*t*) was calculated by the following equation,

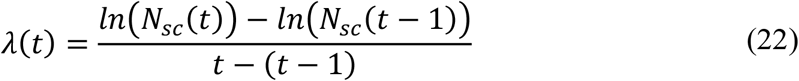

### Neuroprotective strategies

Pharmacological or surgical therapies that abolish the pathological oscillations in STN or block the receptors on SNc can be neuroprotective and might slow down the progression of SNc cell loss (Rodriguez et al., 1998).

#### Glutamate inhibition therapy

Glutamate drug therapy can have neuroprotective effect on SNc in two ways – 1) Inactivation of NMDA (N-methyl-D-aspartate), AMPA (2-amino-3-(5-methyl-3-oxo-1, 2-oxazole-4-yl) propanoic acid) or excitatory metabotropic glutamate (Group-I – mGluR1/5) receptors (mGluR) by glutamate antagonists, and 2) Activation of metabotropic glutamate (Group-II/III – mGluR2,3/4,6,7,8) receptors by glutamate agonists. NMDA antagonist MK-801 showed reduction of SNc cell loss in the neurotoxic rats (Turski et al., 1991; Zuddas et al., 1992b; Brouillet and Beal, 1993; Blandini et al., 2001; Armentero et al., 2006) and primates (Zuddas et al., 1992a, 1992b). AMPA antagonist such as NBQX (Merino et al., 1999), LY-503430 (Murray et al. 2003) and LY-404187 (O’Neill et al., 2004) exhibited neuroprotection of SNc cells in the neurotoxic animal models. mGluR-5 antagonist MPEP and MTEP showed neuroprotection in 6-OHDA lesioned rats (Armentero et al., 2006; Hsieh et al., 2012; Ferrigno et al., 2015; Fuzzati-Armentero et al., 2015) and MPTP-treated primates (Masilamoni et al., 2011) respectively. Broad-spectrum group II (Battaglia et al. 2003; Murray et al. 2002; Vernon et al. 2005) and group III (Vernon et al., 2005; Austin et al., 2010) agonists showed neuroprotection in neurotoxic rats. Selective mGluR2/3 agonist 2R,4R APDC (Chan et al., 2010) and mGluR4 agonist VU0155041 (Betts et al., 2012) significantly attenuated SNc cell loss in 6-OHDA lesioned rats.

The glutamate drug therapy was implemented in the proposed excitotoxicity model by the following criterion,

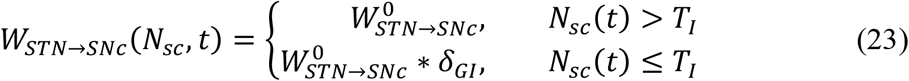

where, *W*_*STN*→*SNc*_(*N_sc_*, *t*) is the instantaneous change in synaptic weight of STN to SNc based on the number of surviving SNc neurons at time (*t*) (*N_sc_*(*t*)), 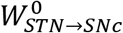 is the basal connection strength of STN to SNc, *δ_GI_* is the proportion of glutamate inhibition, *T_I_* is the percentage of SNc cell loss 〈25|50|75〉 at which therapeutic intervention was employed. In the present study, we have considered 25%, 50% and 75% SNc cell loss as early, intermediate and late stages of disease progression respectively.

#### Dopamine restoration therapy

The neuroprotective effects of DA agonists therapy are thought to be due to one or more of the following mechanisms: 1) L-DOPA sparing, 2) Autoreceptor effects, 3) Antioxidant effects, 4) Antiapoptotic effects and 5) Amelioration of STN-mediated excitotoxicity (Grandas 2000; Olanow, Jenner, and Brooks 1998; Schapira 2003; Zhang and Tan 2016). In the present study, we focus on amelioration of STN-mediated excitotoxicity. DA agonists can restore the dopaminergic tone in the dopamine-denervated brain, which results in increased inhibition in STN, thereby diminishing STN-induced excitotoxicity in SNc neurons (Olanow et al., 1998; Schapira and Olanow, 2003; Piccini and Pavese, 2006; Vaarmann et al., 2013).

The dopamine agonist therapy was implemented in the proposed excitotoxicity model by the following criterion,

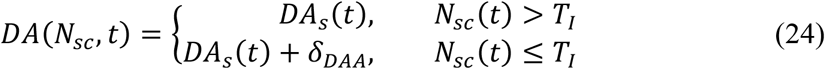

where, *DA*(*N_sc_*,*t*) is the instantaneous change in dopamine level based on the number of surviving SNc neurons at time (*t*) (*N_sc_*(*t*)), *DA_s_*(*t*) is the instantaneous dopamine signal from the SNc neurons, *δ_DAA_* is the proportion of dopamine content restoration, *T_I_* is the percentage of SNc cell loss 〈25|50|75〉 at which therapeutic intervention was employed.

#### Subthalamotomy

Subthalamotomy is still quite a prevalent treatment amongst patients in advanced stages of PD where patients stop responding to L-DOPA (wearing-off) or chronic L-DOPA therapy results in motor complications such as L-DOPA Induced Dyskinesias (LID) (Alvarez et al., 2009; Obeso et al., 2017). It was reported that STN lesioning exhibits neuroprotective effect which acts as an antiglutamatergic effect in neurotoxic animal models (Piallat et al., 1996; Chen et al., 2000; Carvalho and Nikkhah, 2001; Paul et al., 2004; Wallace et al., 2007; Jourdain et al., 2014).

STN ablation was implemented in the proposed excitotoxicity model by the following criterion,

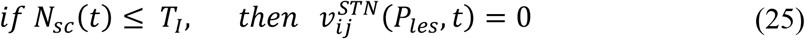

where, *P_les_* is the lesion percentage of STN which is selected from the following range: {5,10, 20,40, 60, 80,100}, *N_sc_*(*t*) is the instantaneous number of surviving SNc neurons, *T_I_* is the percentage of SNc cell loss 〈25|50|75〉 at which therapeutic intervention was employed.

#### Deep brain stimulation (DBS) in STN

DBS therapy is preferred over ablation therapy of STN due to the potentially irreversible damage to the stimulated brain region in ablation therapy. It has been reported that long-term stimulation (DBS) of STN results in slowdown of the progression of SNc cell loss in animal models (Benazzouz et al., 2000; Maesawa et al., 2004; Temel et al., 2006; Wallace et al., 2007; Spieles-Engemann et al., 2010; Musacchio et al., 2017).

The DBS electrical stimulation was given in the form of current or voltage pulses to the target neuronal tissue (Cogan, 2008). The effect of DBS therapy was modeled as external stimulation current given to the entire or part of STN module in the form of Gaussian distribution (Rubin and Terman, 2004; Hauptmann and Tass, 2007; Foutz and McIntyre, 2010; Mandali and Chakravarthy, 2016). The DBS parameters such as amplitude (*A_DBS_*), frequency 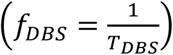 and pulse width 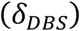 were adjusted by using clinical settings as a constraint (Moro et al., 2002; Garcia et al., 2005), in order to reduce the synchrony in STN population along with the minimal rise in the firing rate. In addition to exploring DBS parameters, a range of stimulus waveforms (such as rectangular monophasic and biphasic current pulses) and different types of stimulation configurations (such as single contact point (SCP), four contact points (FCP) and multiple contact points (MCP)) were also implemented (fig. 2) (Cogan, 2008; Lee et al., 2016).

**Figure 2:**
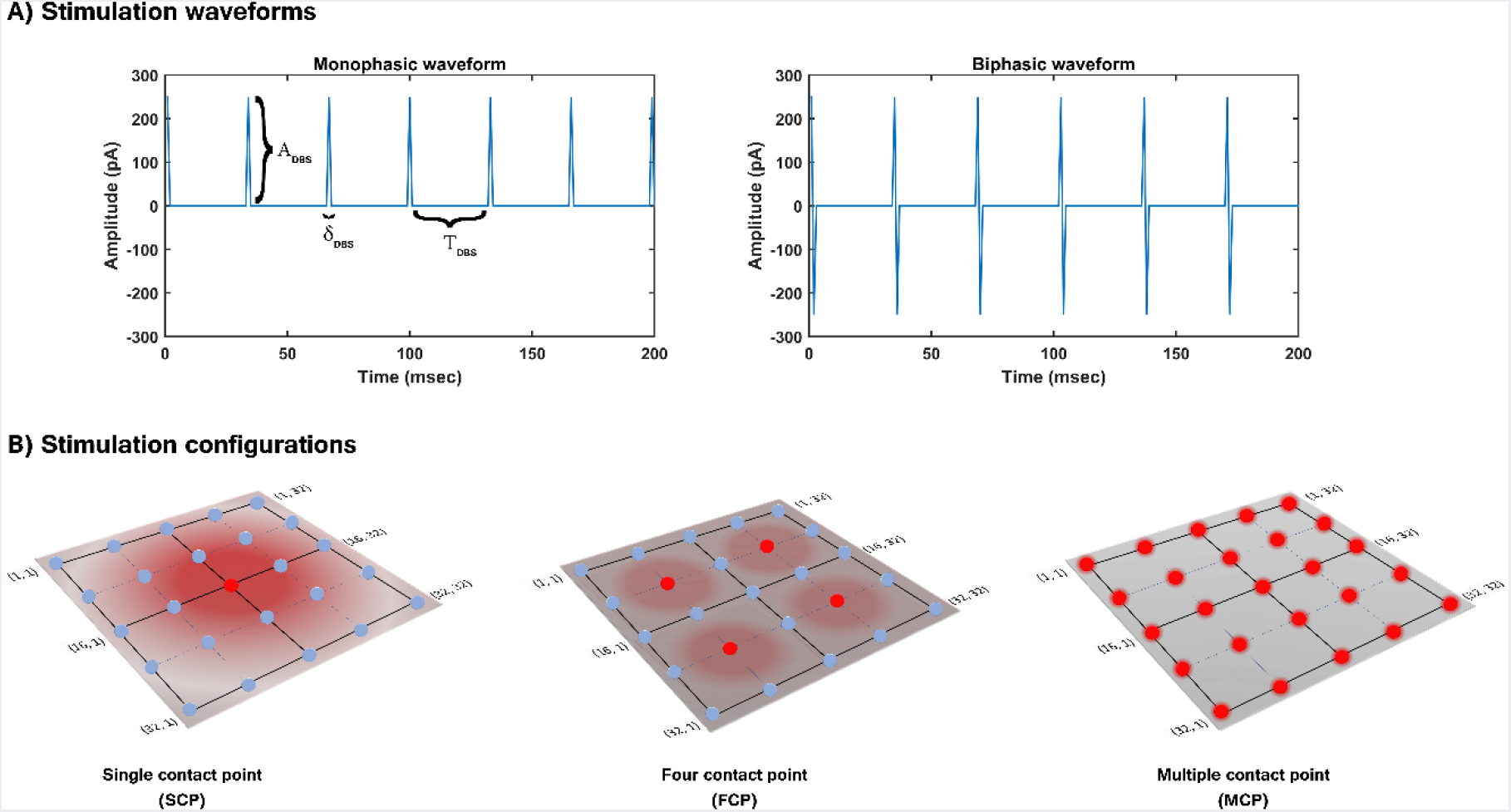
(A) DBS stimulation waveforms (B) DBS stimulation configurations.

In the present study, the current pulses which given to neuronal network are in the form of monophasic and biphasic waveforms. The monophasic current pulse (*P_MP_*) was generated as the following,

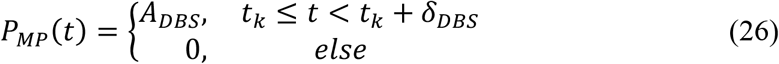

where, *t_k_* are the onset times of the current pulses, *A_DBS_* is the amplitude of the current pulse, *δ_DB_s* is the current pulse width.

The biphasic current pulse (*P_BP_*) was generated as the following,

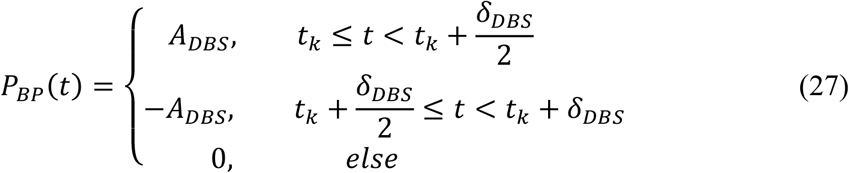

where, *t_k_* are the onset times of the current pulses, *A_DBS_* is the amplitude of the current pulse, *δ_DBS_* is the current pulse width.

The influence of stimulation on a particular neuron will depend on the position of the stimulation electrode in the neuronal network (Cogan, 2008). The effect of stimulation will decay as the distance between electrode position (*i_c_*,*j_c_*) and neuronal position (*i,j*) increased which was modeled as a Gaussian neighbourhood (Mandali et al., 2016). We have assumed that the center of the electrode to be the mean of the Gaussian which coincides with the lattice position (*i_c_*,*j_c_*) and the spread of stimulus current was controlled by the width of the Gaussian (*σ*)

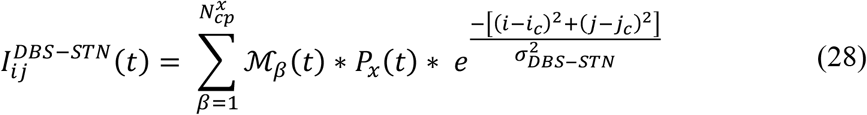

where, 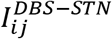 is the DBS current received by STN neuron at position (*i,j*) considering lattice position (*i_c_*,*j_c_*) as the electrode contact point at time (*t*), *ℳ_β_*(*t*) is the indicator function which controls the activation of stimulation site 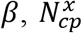 is the number of activated stimulation contact points for different stimulation configurations *x* = {*SCP,FCP,MCP*} 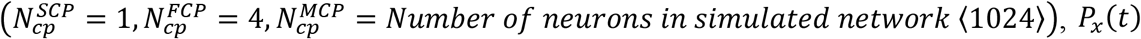, *P_x_*(*t*) is the current pulse at time *t* for *x* = {*MP, BP*}, *σ_DBS-STN_* is used to control the spread of stimulus current in STN network.

DBS was implemented in the proposed excitotoxicity model by the following criterion,

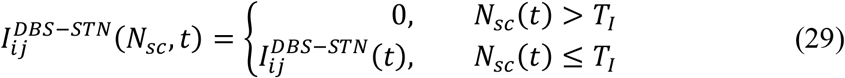

where, 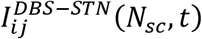 is the instantaneous change in the stimulation current to STN neuron at position (*i,j*) based on the number of surviving SNc neurons at time (*t*)

#### Antidromic activation

The mechanism of how DBS alleviates advanced PD symptoms is not clear. One of the theories behind therapeutic effect of DBS is activation of afferent connections of STN which results in antidromic activation of cortical, GPi or GPe neurons (Chiken and Nambu 2016; Hammond et al. 2008; Kang and Lowery 2014; Lee et al. 2004; McIntyre et al. 2004; Montgomery and Gale 2008; Oluigbo, Salma, and Rezai 2015). In our study, we implemented the antidromic activation of GPe during DBS therapy. Antidromic activation was implemented similarly to (Mandali et al., 2016), where a percentage of DBS current given to STN neurons were given directly to GPe neurons. Similar to DBS applied to STN, external stimulation current was given to GPe neuron in the form of Gaussian distribution. The specifications of antidromic activation were described by the following equation,

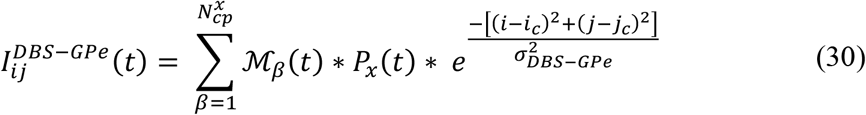

where, 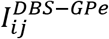 is the DBS current received by GPe neuron at position (*i,j*) considering lattice position (*ic,jc*) as the electrode contact point, *ℳ_β_*(*t*) is the indicator function which controls the activation of stimulation site *β*, 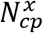 is the number of activated stimulation contact points for different stimulation configurations *x* = {*SCP,FCP,MCP*} 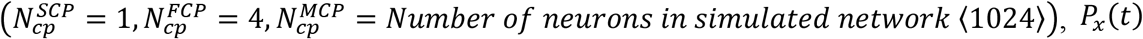, *P_x_*(*t*) is the current pulse at time *t* for *x* = {*MP,BP*}, *A_DBS-GPe_* is the portion of DBS current pulse amplitude given as antidromic activation to GPe neurons (*pA*), σ*_DBS-GPe_* is used to control the spread of stimulus current in GPe ensemble.

The DBS therapy with antidromic activation was implemented in the proposed excitotoxicity model by the following criterion,

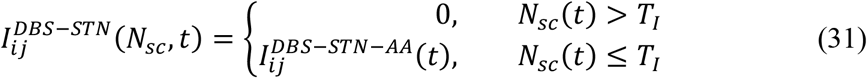

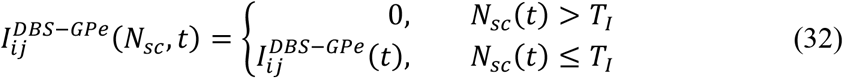

where, 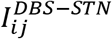 is the DBS current received by STN neuron at position (*i,j*) considering lattice position (*ic,jc*) as the electrode contact point with antidromic activation 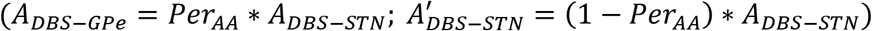, *Per_AA_* is the portion of *A_DBS–STN_* applied as 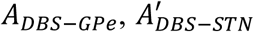 is the portion of DBS current pulse amplitude given to STN neurons during antidromic activation (*pA*)

#### STN axonal & synaptic failures

It has been reported that during continuous high-frequency stimulation of the STN result in synaptic depression in the STN as observed from *in vitro* recordings (Ledonne et al., 2012). The synaptic depression caused by increased STN activity during DBS arises due to an amalgamation of axonal and synaptic failures in the STN (Ammari et al. 2011; Carron et al. 2013; Moran et al. 2011, 2012; Oluigbo, Salma, and Rezai 2015; Rosenbaum et al. 2014; Shen and Johnson 2008; Zheng et al. 2011).

The effect of synaptic depression due to DBS of the STN was implemented by the following criterion,

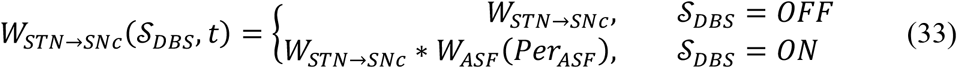

where, 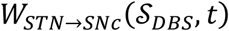 is the instantaneous change in synaptic weight of STN to SNc based 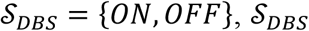 is DBS stimulation, *W_ASF_* is the weight matrix based on the percentage of axonal and synaptic failures (*Per_ASF_*)

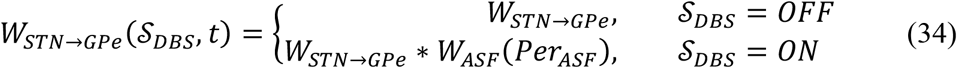

where, 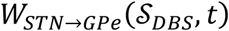 is the change in synaptic weight of STN to GPe based 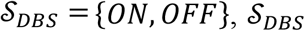 is DBS stimulation, *W_ASF_* is the weight matrix based on the percentage of axonal and synaptic failures (*Per_ASF_*)

### Network analysis

For the analysis of network dynamics, we have used Multiple neural spike train data analysis procedures (Brown et al., 2004) such as peri-stimulus time histogram (Bretschneider and De Weille, 2006), network synchrony (Pinsky and Rinzel, 1995) and burst index measure (van Elburg and van Ooyen, 2004).

The proposed model was tuned and simulated for normal and PD conditions characterized by loss of SNc cells. Under normal conditions, the loop interactions between SNc and STN are such that, the stress levels in SNc do not exceed the apoptotic threshold, and therefore the SNc cells survive. But if a critical number of SNc cells die, the reduced SNc size leads to disinhibition of STN, which becomes overactive, due to which SNc cells receive over-excitation leading to neurodegeneration. Thus, the initial loss of SNc cells leads to a runaway effect, leading to an uncontrolled loss of cells in the SNc, characterizing the underlying neurodegeneration of PD.

## III. Results

We have investigated the Izhikevich parameters of STN, SNc and GPe which were chosen from the literature (Michmizos and Nikita, 2011; Cullen and Wong-Lin, 2015; Mandali et al., 2015) for their characteristics firing pattern and other biological properties (Figure-3). We then extensively studied the effect of lateral connections in the network of neurons (Figure-4, 6, 7) and compared with experimental data (Figure-5). Next, we have explored the effect of dopamine on the network of GPe, SNc and STN neurons and compared with published data (Figure-8, 9, 10).

Then, we showed the results of the proposed excitotoxicity model which exhibits STN-mediated excitotoxicity in SNc (Figure-11, 12, 13, 14) and studied their sensitivity to parameter uncertainty (Figure-15, 16). Finally, we have explored current therapeutics such as glutamate inhibition (Figure-17, 18, 19), dopamine restoration (Figure-20, 21, 22), subthalamotomy (Figure-23, 24, 25) and deep brain stimulation (Figure-26, 27, 28, 29, 30) which might have neuroprotective effect on the progression of SNc cell loss.

### Characteristics firing of different neuronal types

The behavior of single neuron models of the three different neuronal types involved in the excitotoxicity model for different external current input can be seen in fig. 3. In the proposed model, we adjusted 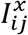 and other parameters of the Izhikevich model such that the basal firing frequencies of the different neuronal types match with experimental data (Modolo et al., 2007; Tripathy et al., 2014, 2015). The adjusted values can be seen in the *Table-1*.

**Figure 3:**
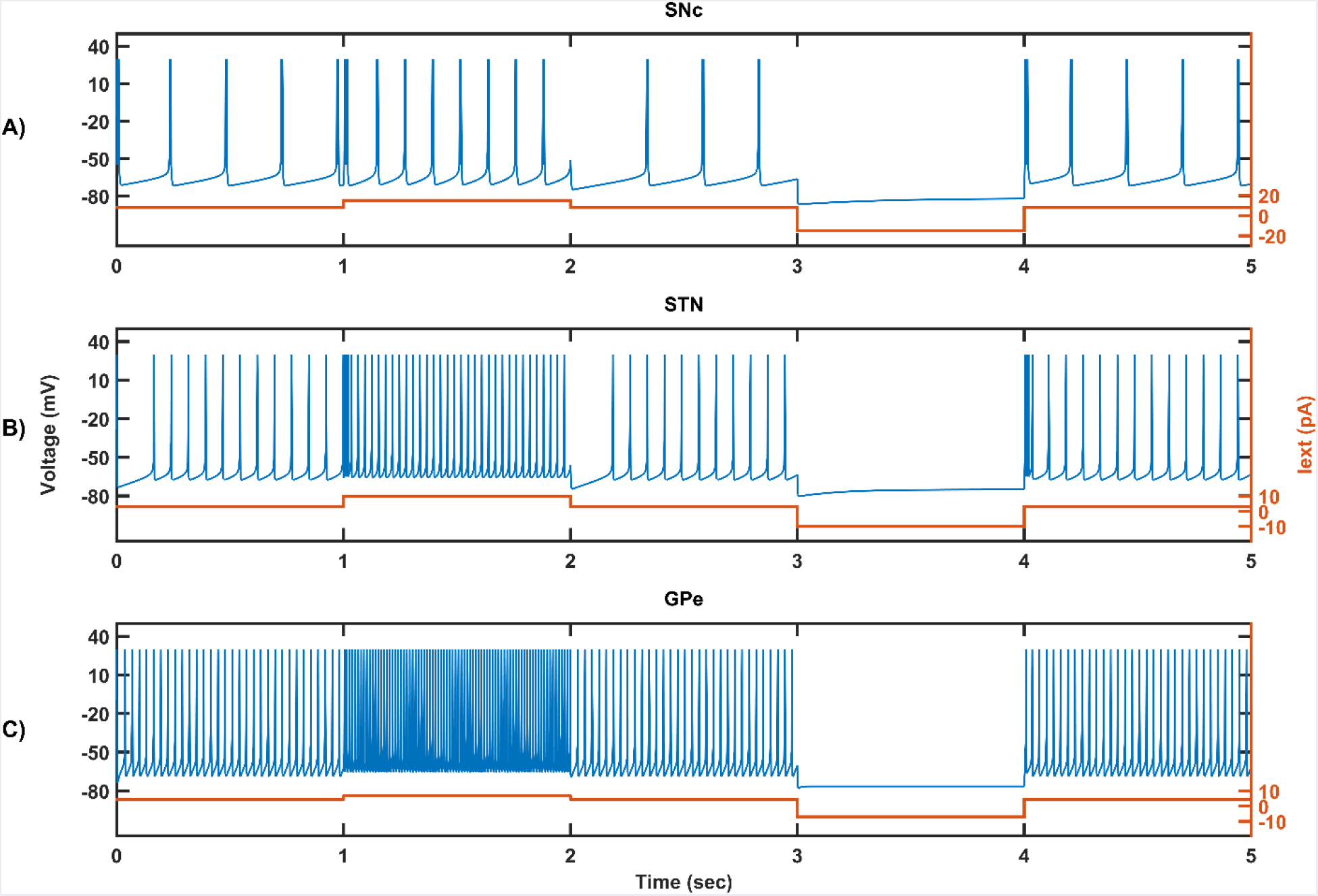
Firing characteristics of SNc (A), STN (B) and GPe (C) varying external currents (orange line).

The SNc neurons experimentally exhibit two distinct firing patterns: low-frequency irregular tonic or background firing (3-8 *Hz*) and high-frequency regular phasic or burst firing (~20 *Hz*) (Grace and Bunney, 1984a, 1984b). The Izhikevich parameters which were chosen for SNc neurons were able exhibits both type of firing patterns. Other properties such as doublet-spikes which were occasionally observed experimentally (Grace and Bunney, 1983) were also exhibited (fig. 3A). In the present model, SNc neurons basal firing rate were required to be ~4 Hz which is in the range of 3-8 Hz observed experimentally (Grace and Bunney, 1984a). Similar to SNc, STN neurons also exhibits tonic pacemaking firing and phasic high-frequency bursting (Beurrier et al., 1999; Allers et al., 2003). The basal firing rate of STN neurons were required to be ~13 Hz which is in the range of 6-30 Hz observed experimentally (Allers et al., 2003; Lindahl et al., 2013). The STN neurons also exhibits characteristic inhibitory rebound which was observed experimentally (fig. 3B) (Hamani et al., 2004; Johnson, 2008). Unlike SNc and STN, GPe neurons exhibits tonic high-frequency firing which was interpreted by bursts and pauses (Kita and Kita, 2011; Hegeman et al., 2016). The Izhikevich parameters which were chosen for GPe neurons were able to exhibits high-frequency firing without any bursts. The basal firing rate of GPe neurons were required to be ~30 Hz which is in the range of 17-52 Hz observed experimentally (Lindahl et al., 2013).

### Network activity of all three neuronal types

#### Behavior regimes with varying collaterals strength and radius

We now study the network dynamics of each of the three neuronal types in a 2D array with lateral connections. The effect of network properties like the strength and neighborhood size of the lateral connections on the firing properties like average firing rate, synchrony, and burst index is studied (fig. 4). The suitable values of lateral connections strength and radius for each neuronal type were chosen in correlation with experimental data (Humphries et al., 2006; Tepper and Lee, 2007). The selected values can be seen in the *Table-1*.

**Figure 4:**
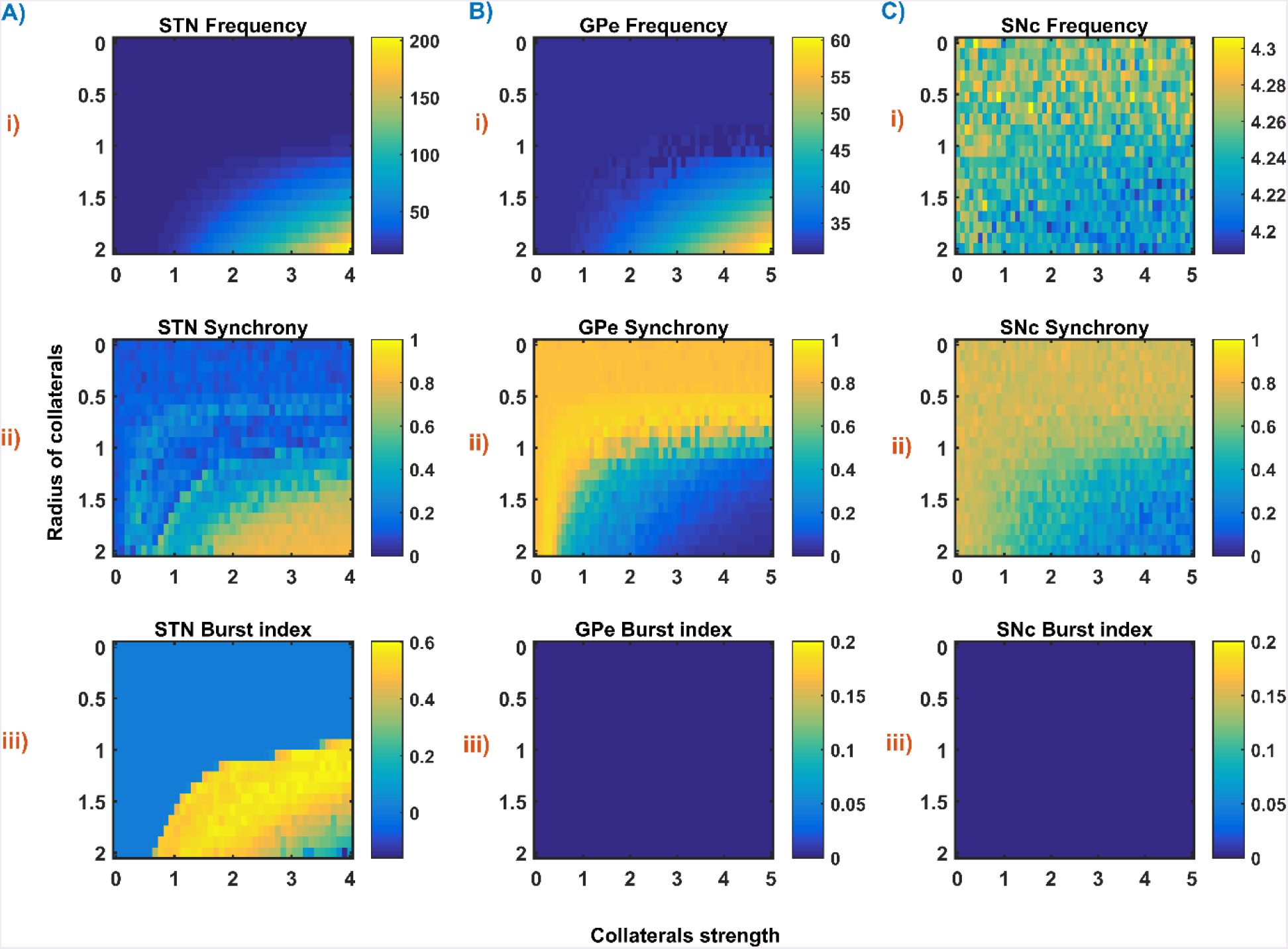
The response of STN (A), GPe (B) and SNc (C) populations with varying lateral strength and radius at the level of network properties (Frequency (i), Synchrony (ii) and Burst index (iii))

#### Basal population activity

As specified above, 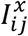, *A_x_* and *R_x_* was adjusted such that the basal population activity correlated well with the experimental data (Humphries et al., 2006; Tepper and Lee, 2007) as shown in the fig. 5.

**Figure 5:**
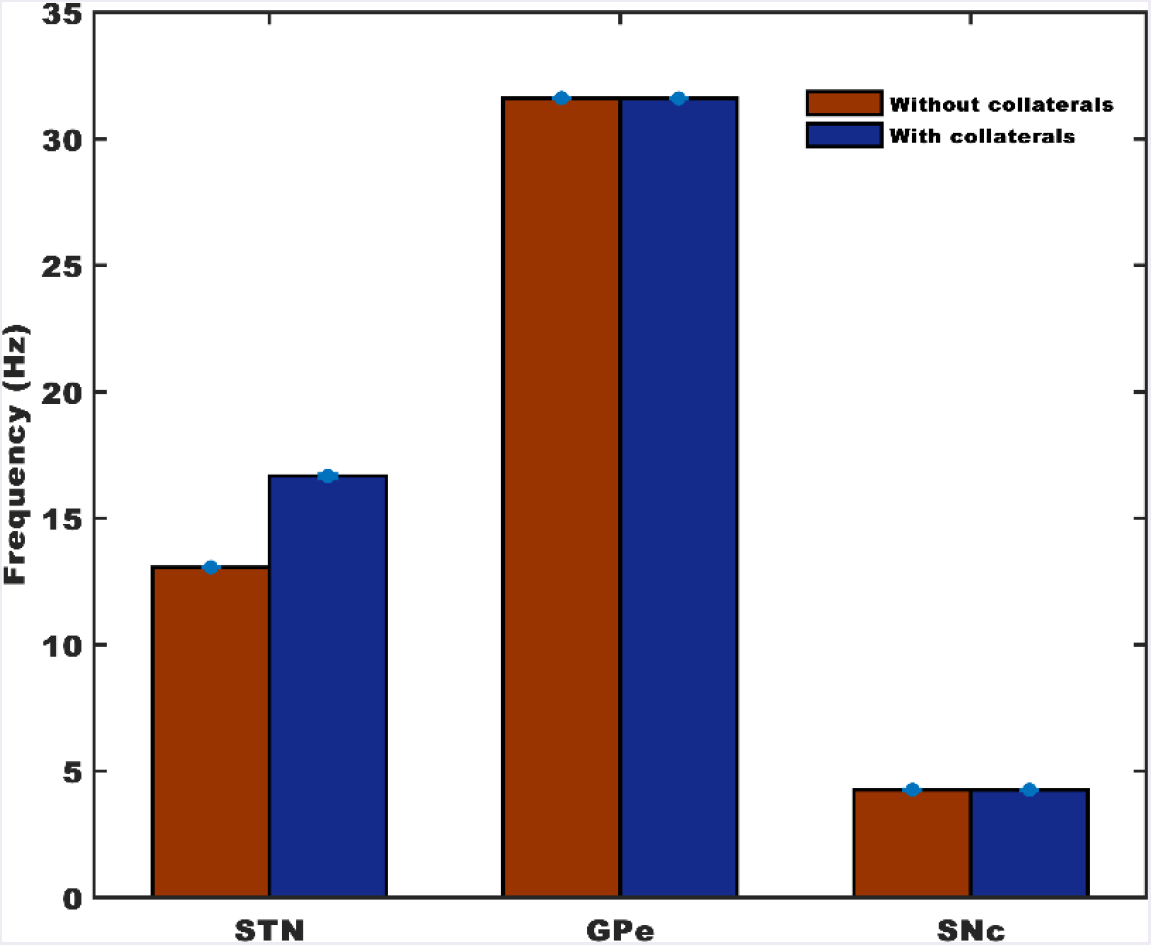
Basal population activity of each neuronal type with and without collaterals.

#### STN population activity

The network dynamics of STN plays a vital role in the proposed model of excitotoxicity, in this scenario we have studied the role of lateral connections in regulating STN network properties. The basal STN population activity without lateral connections showed normal spiking without any bursting type of behavior as seen in the fig. 6. The fast Fourier transform (FFT) analysis showed that the frequency content in the STN population exhibits non-bursting activity as seen in the fig. 4(inset). Contrarily, the basal STN population activity with lateral connections showed the bursting type of activity as seen in the fig. 7. FFT analysis also showed that the frequency content in STN population exhibits bursting activity as seen in the fig. 7(inset).

**Figure 6:**
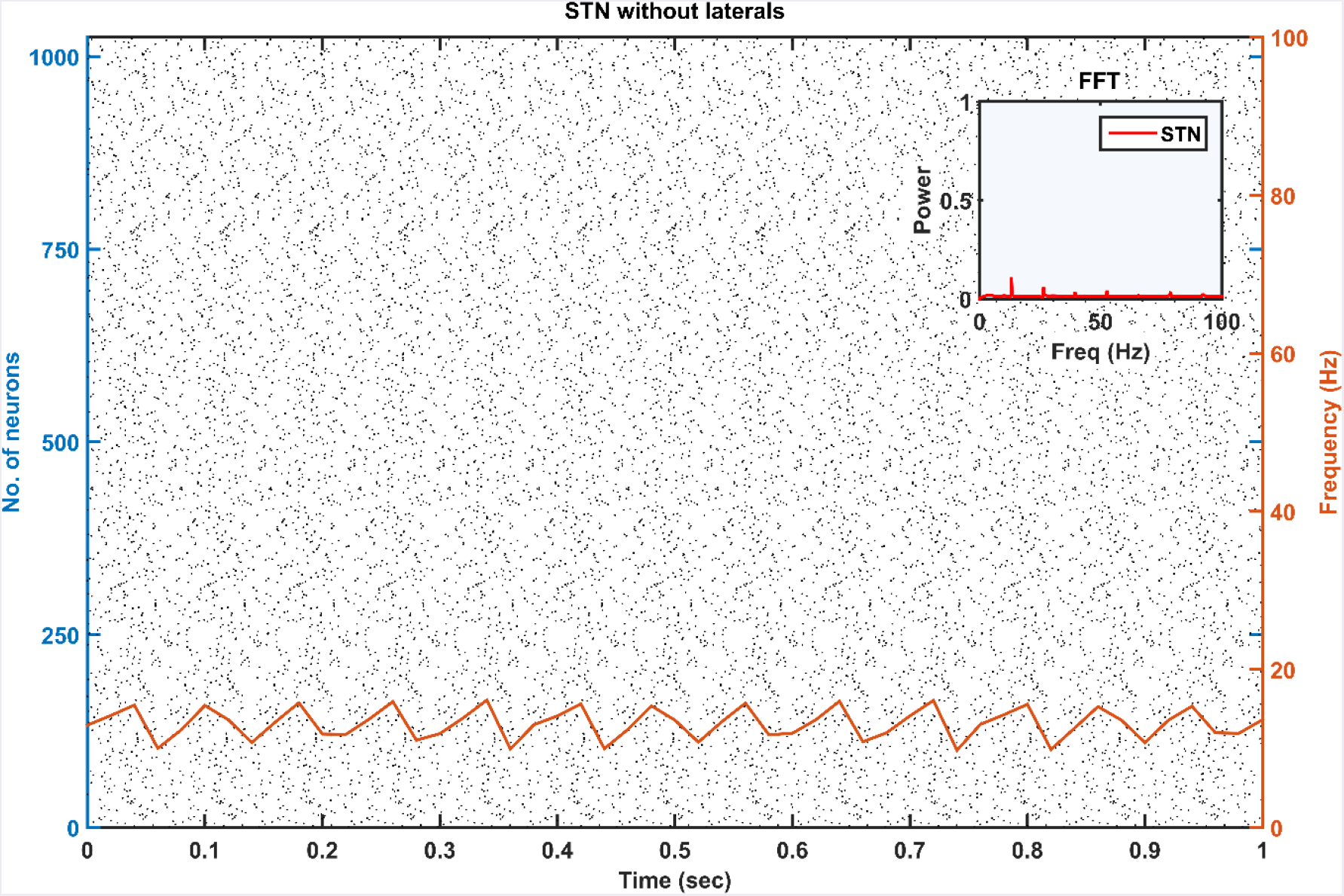
Raster plot of basal population activity of STN without collaterals are overlaid with spiking-count firing rate (orange line); Inset – Frequency content in STN population.

**Figure 7:**
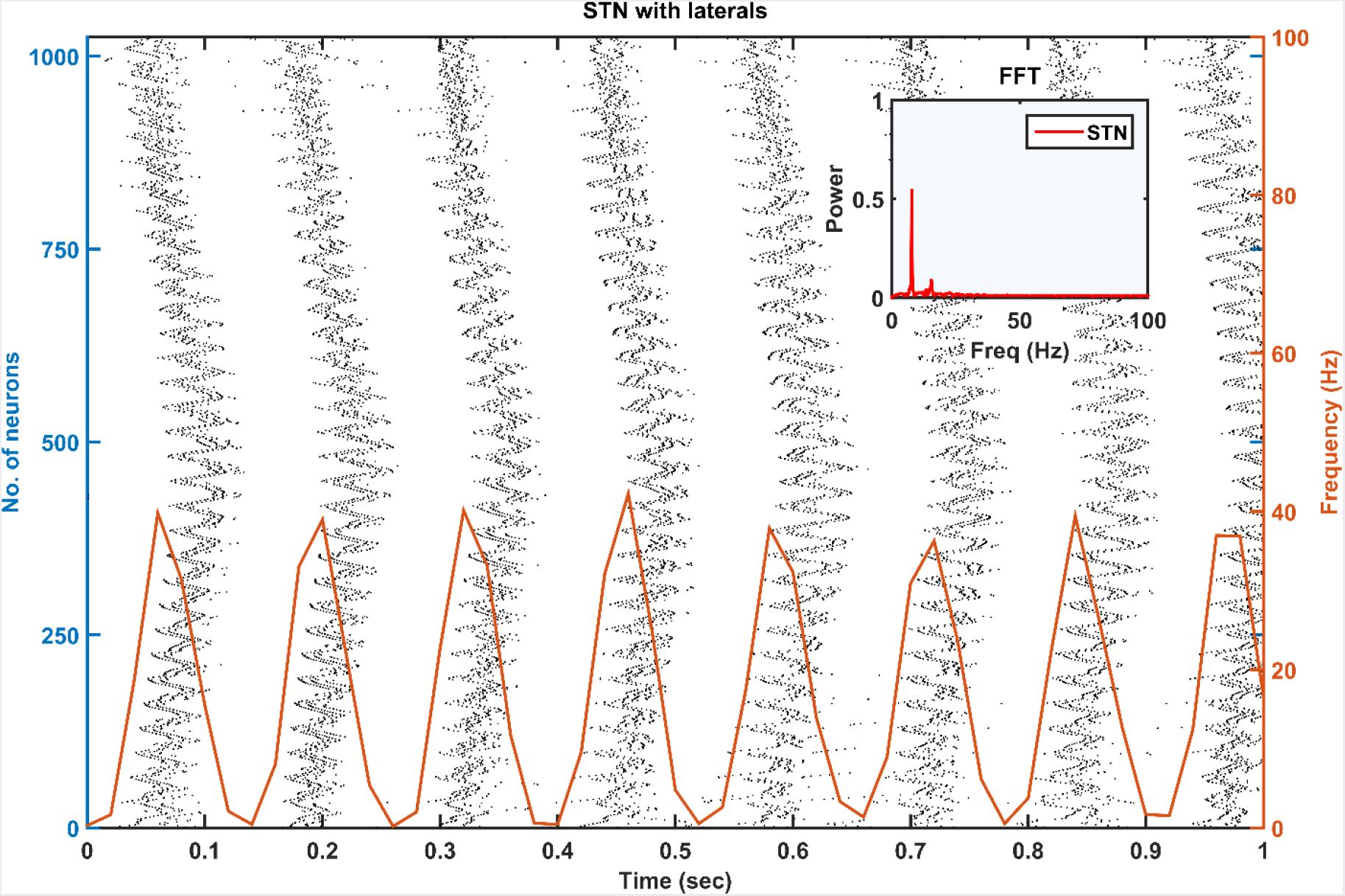
Raster plot of basal population activity of STN with collaterals are overlaid with spike-count firing rate (orange line); Inset – Frequency content in STN population.

### Dopamine effect on the basal activity

#### Individual populations

From the simulated results, it is clear that as DA level increases the mean firing rate decreases in STN, increases in GPe and decreases in SNc as shown in the fig. 6. The network synchrony decreases in all neuronal populations as DA levels increases. But in the case of STN, the decrease is not monotonic as can be seen in the fig. 8A(ii) where high synchrony was observed at moderate levels of DA, with synchrony falling on either side. This high synchronicity at moderate levels of DA is a result of change in firing pattern from asynchronous bursting to synchronous spiking which can be correlated with burst index (see the fig. 8A(iii)) in STN population. In the dopamine-depleted condition, STN shows the bursting type of firing pattern which was exhibited by our model consistent with published studies (Vila et al., 2000; Ammari et al., 2011; Park et al., 2015). The following trend of STN activity was observed when DA level increases from 0 to 1: synchronous bursting, asynchronous bursting, synchronous spiking and asynchronous spiking. At very low DA levels (0-0.1), the STN exhibits regular bursting (fig. 8A(iii)) with high synchrony (fig. 8A(ii)). At low DA levels (0.1-0.3), the STN exhibits an irregular mixed mode of bursting and singlet-spiking with low synchrony (fig. 8A(ii)). At moderate DA levels (0.3-0.7), the STN exhibits regular singlet-spiking (fig. 8A(iii)) with high synchrony (fig. 8A(ii)). And at high DA levels (0.3-1), the STN exhibits irregular singlet-spiking with low synchrony (fig. 68(ii)).

**Figure 8:**
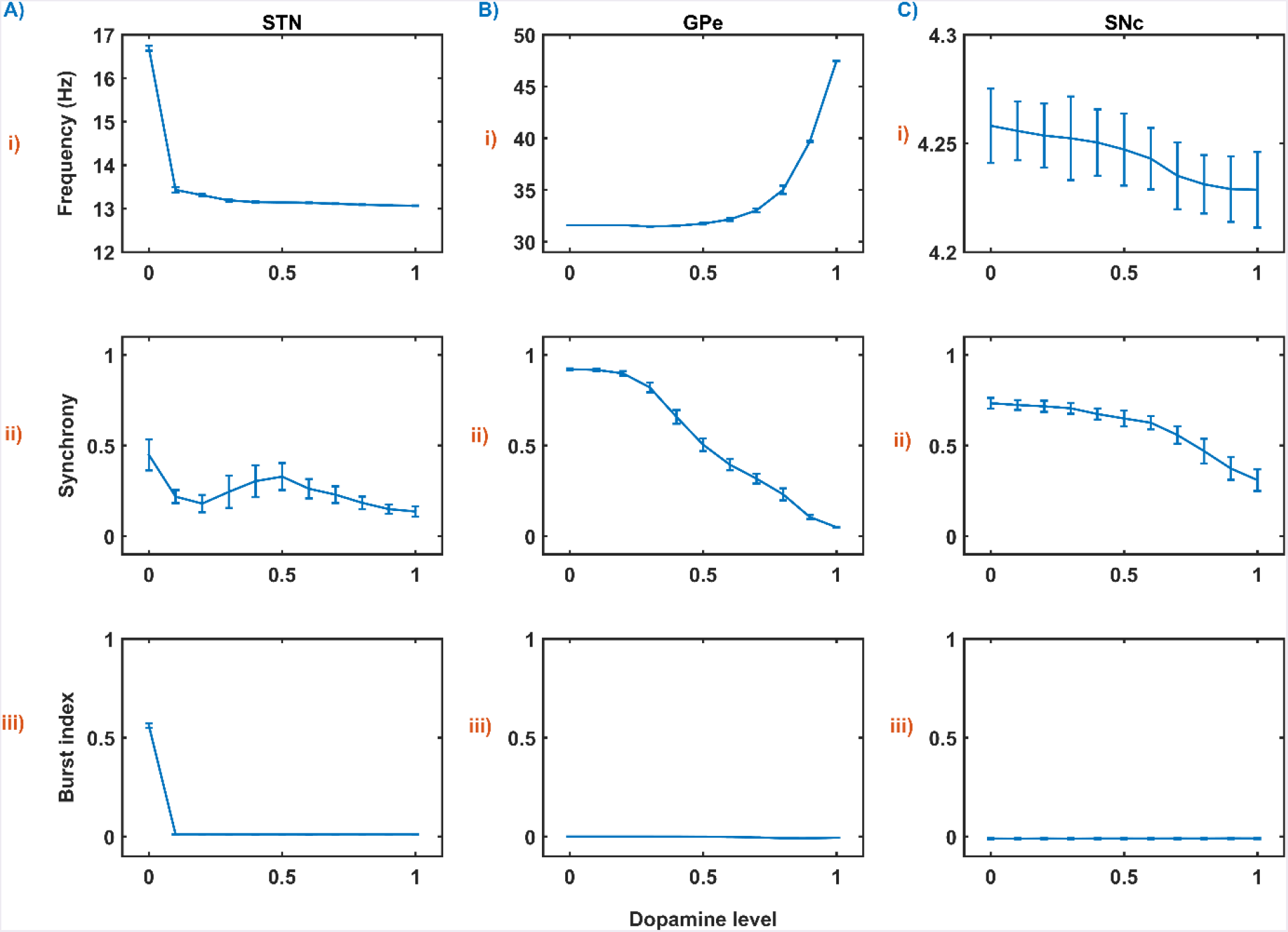
The response of STN (A), GPe (B) and SNc (C) populations with varying dopamine levels at the level of network properties (Frequency (i), Synchrony (ii) and Burst index (iii)).

#### STN-GPe network

STN-GPe dynamics is known to play an important role in PD pathological oscillations that are thought to be strongly related to the cardinal symptoms of PD (Bergman et al., 1994; Brown, 2003; Litvak et al., 2011; Park et al., 2011). Numerous computational models were developed to explain the pathological oscillations in STN-GPe (Terman et al., 2002; Pavlides et al., 2015; Shouno et al., 2017). The connectivity pattern between STN and GPe was explored by using a conductance-based model (Terman et al., 2002) which exhibited different rhythmic behaviors. In our model, the connectivity pattern between STN and GPe was considered to be dopamine-dependent (Cragg et al., 2004; Mandali et al., 2015) and spontaneous activity of STN-GPe network was studied with no external input current. Under normal DA conditions, low synchrony and minimal oscillations were exhibited by the STN-GPe network (fig. 9B) (Kang and Lowery, 2013). But in dopamine-depleted conditions, high synchrony and higher rate of oscillations were exhibited in STN-GPe network as observed in previous studies (fig. 9A) (Brown et al., 2001; Park et al., 2011; Lintas et al., 2012; Kang and Lowery, 2013). From simulated results, beta range oscillations were observed in STN population under dopamine-depleted conditions as observed in previous studies (Brown et al., 2001; Park et al., 2010; Pavlides et al., 2015).

**Figure 9:**
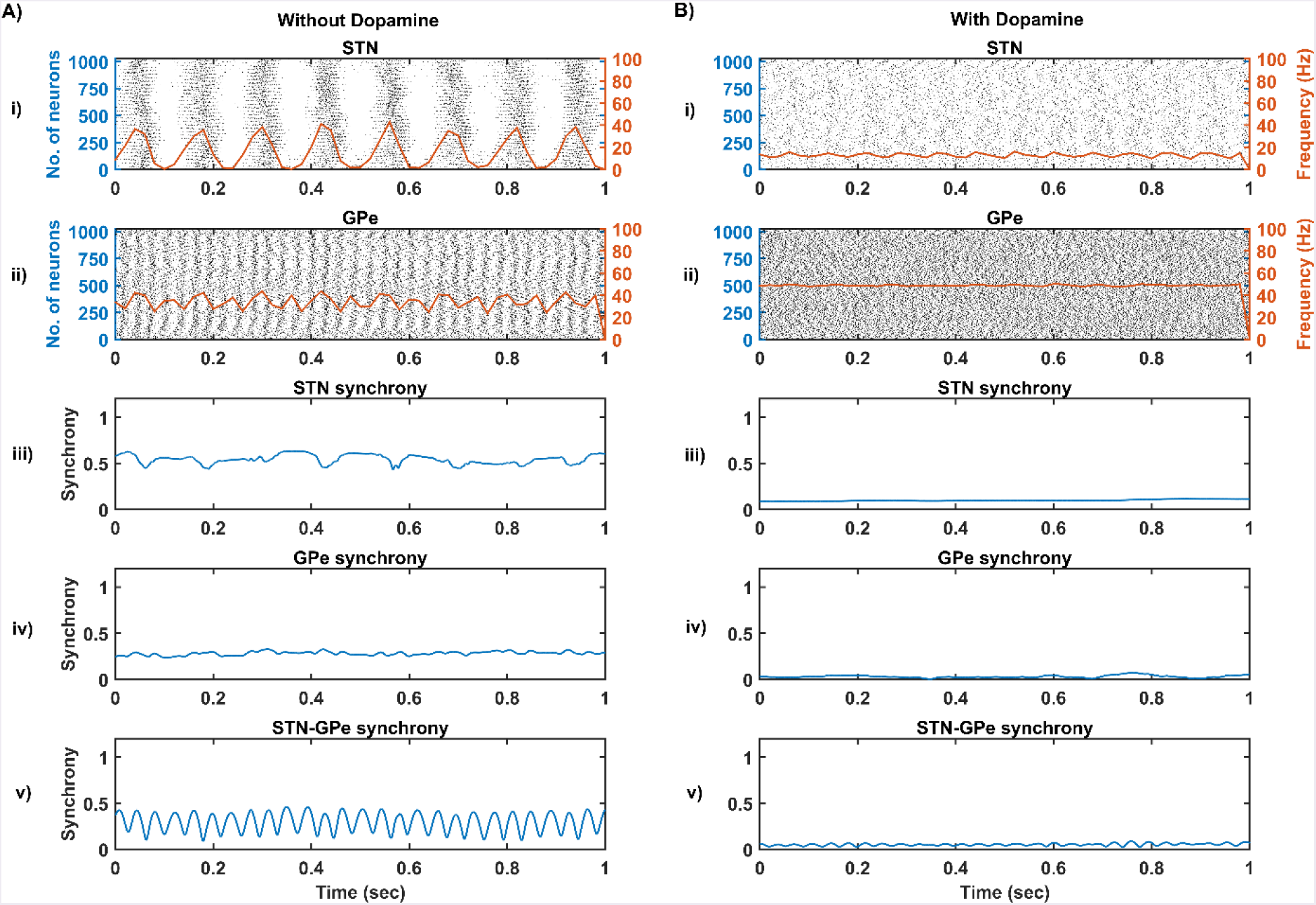
The response of STN-GPe network without (A) & with (B) dopamine – Raster plots of STN (i) & GPe (ii) populations overlaid with spike-count firing rate (orange line), Synchrony plots of STN (iii), GPe (iv) and combined STN-GPe (v)

### Influence of SNc dopamine on STN activity

It was reported that dopamine-depleted condition results in pathological oscillations in STN characterized by high synchrony and beta range oscillations (Brown et al., 2001; Weinberger et al., 2006; Park et al., 2010, 2011; Lintas et al., 2012; Pavlides et al., 2015). In our model, the neuronal response of STN with and without SNc projections as shown in the fig. 10. And FFT analysis showed that the frequency content in STN population exhibits bursting as shown in the fig. 10A(iii).

**Figure 10:**
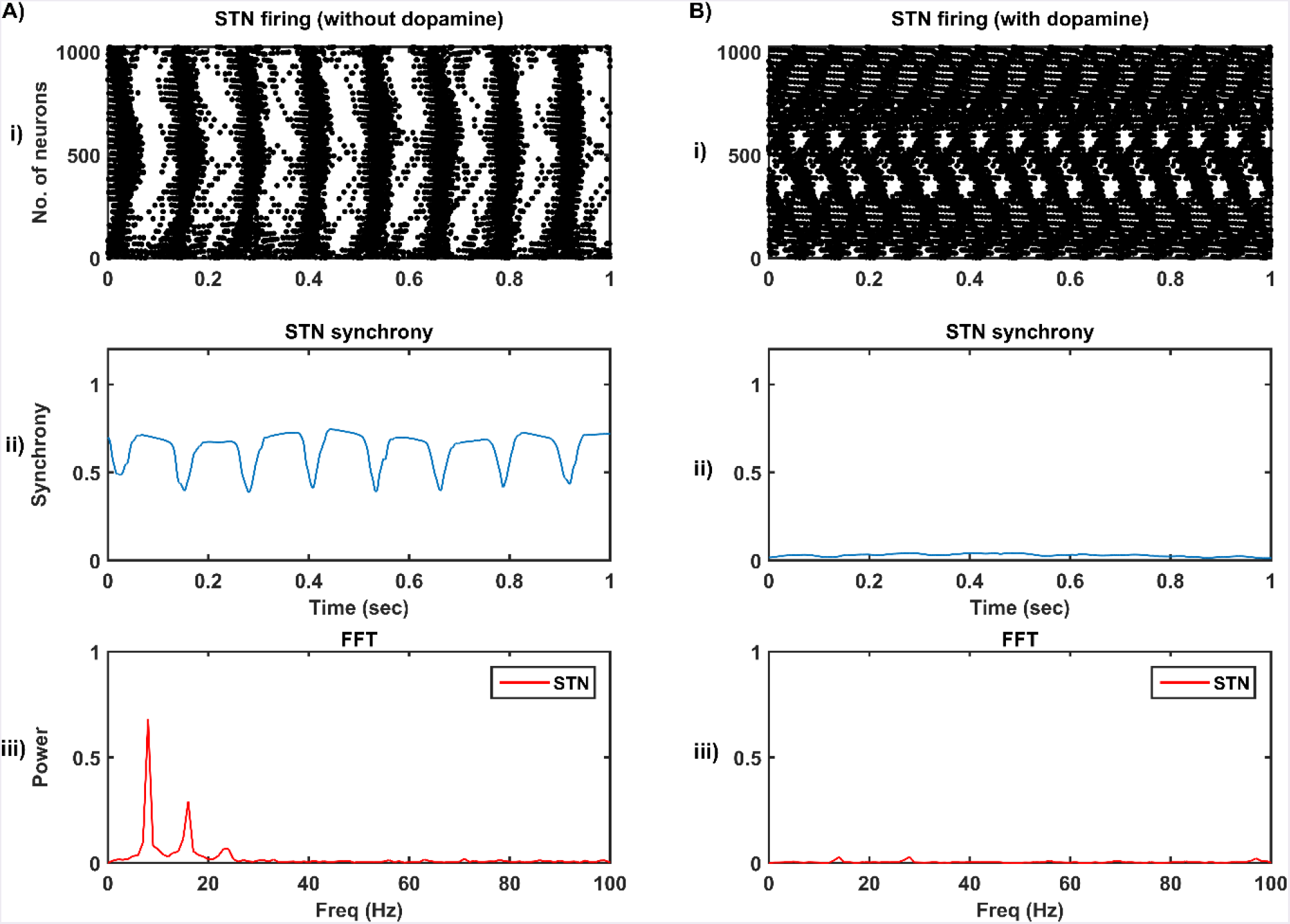
The STN population response (raster plot) without (A) and with (B) SNc projections – network synchrony (ii) and frequency content (iii) in STN.

### STN-induced excitotoxicity in SNc

The proposed excitotoxicity model was able to exhibit STN-mediated excitotoxicity in SNc which was precipitated by energy deficiency (Albin and Greenamyre, 1992; Beal et al., 1993; Greene and Greenamyre, 1996; Rodriguez et al., 1998; Blandini, 2001, 2010; Ambrosi et al., 2014) as shown in the fig. 11. For a more detailed explanation of the excitotoxicity results obtained, we have sub-divided 50 seconds simulation into three parts – I) STN-SNc loop dynamics (normal condition), II) Stress-induced neurodegeneration in SNc (pre-symptomatic PD condition) and III) STN-mediated runaway effect of neurodegeneration in SNc (symptomatic PD condition).

**Figure 11:**
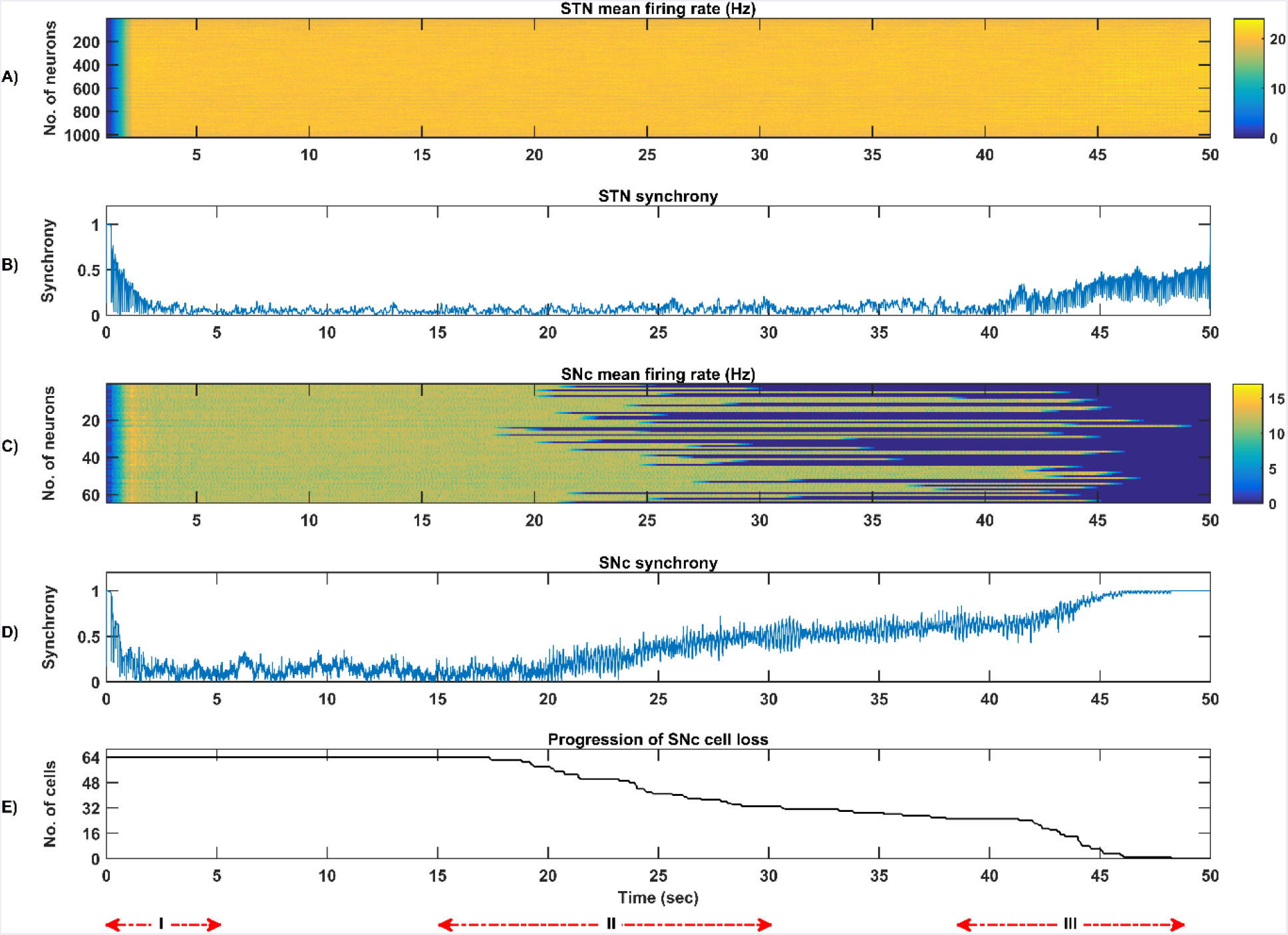
Simulation plots of STN-induced excitotoxicity in SNc – Mean firing rate (1s) of STN (A) & SNc (C), Synchrony of STN (B) & SNc (D), Progression of SNc cell loss (E).

#### (I) STN-SNc loop dynamics

In the first part of the simulation, connectivity between STN and SNc were introduced at *t = 0* and the model exhibited decreased synchrony in STN and SNc over time as a result of short-term plasticity as shown in the fig. 12B. The results showed the pivotal role of dopamine in modulating STN activity (Cragg et al., 2004; Lintas et al., 2012; Yang et al., 2016). The excitatory drive from STN to SNc results in decreased synchrony in SNc due to increased inhibitory drive from lateral connections (fig. 12D). During this whole process, the stress threshold (*S_thres_* = 11.3) was fixed and there was no SNc cell loss due to stress (fig. 12E).

**Figure 12:**
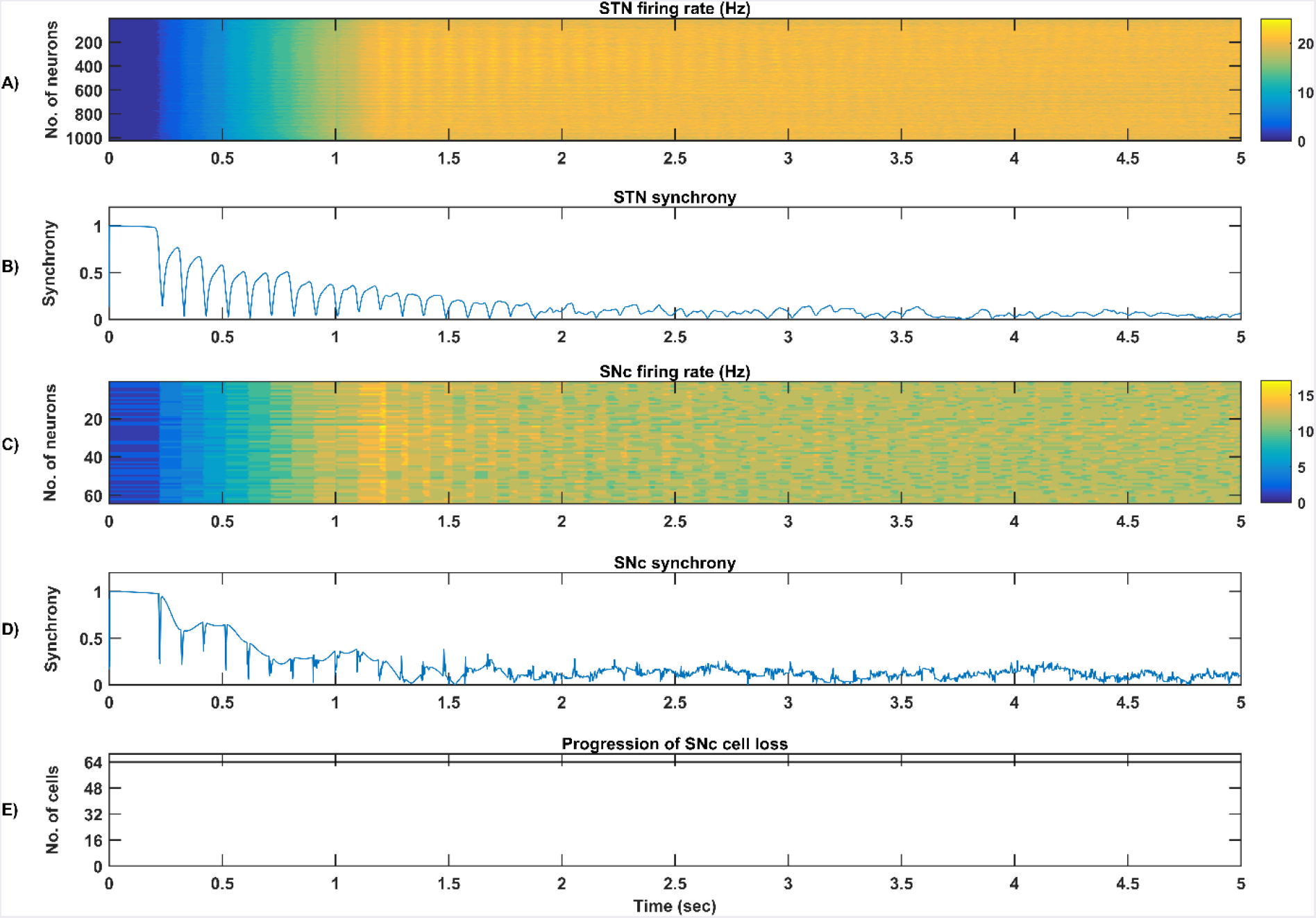
Simulation plots of STN-SNc loop dynamics (Part-I in fig. 11) – Mean firing rate (1s) of STN (A) & SNc (C), Synchrony of STN (B) & SNc (D), Progression of SNc cell loss (E).

#### (II) Stress-induced neurodegeneration in SNc

In the second part of the simulation, stress threshold was slightly reduced from *S_thres_* = 11.3 to *S_thres_* = 10.8 at *t* = 10s to replicate PD-like condition in the model where stress-induced neurodegeneration gets initiated. The model exhibited stress-induced neurodegeneration in SNc where SNc cells start dying when stress variable 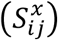 exceeds the stress threshold (*S_thres_*) which acts like an apoptotic threshold (fig. 13E). It was observed that there was no increased synchrony in the STN population as a result of SNc cell loss (fig. 13B). But, there was an increased synchrony in SNc population (fig. 13D) which might be due to reduced inhibitory drive from lateral connections as result of SNc cell loss.

**Figure 13:**
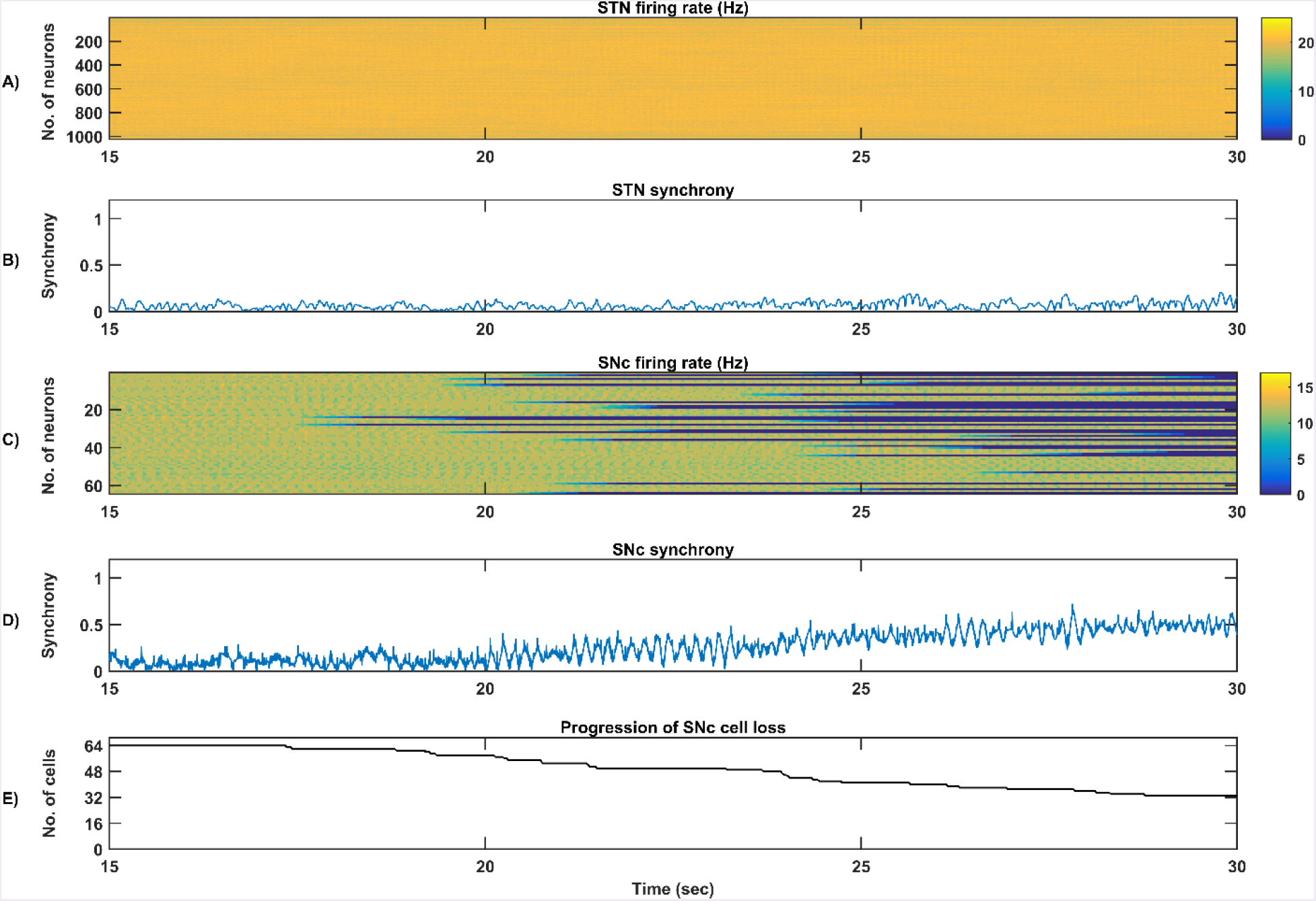
Simulation plots of Stress-induced neurodegeneration in SNc (Part-II in fig. 11) – Mean firing rate (1s) of STN (A) & SNc (C), Synchrony of STN (B) & SNc (D), Progression of SNc cell loss (E).

#### (III) STN-mediated runaway effect of neurodegeneration in SNc

In the third part of the simulation, no parameters were changed but after *t* = 40s there was a rise in STN synchrony as a result of stress-induced SNc cell loss as shown in the fig. 14. After a substantial amount of SNc cell loss (more than 50%) result in increased synchrony (fig. 14B) and firing rates (fig. 14A) in the STN population. As the STN synchrony increased, runaway effect kicks in where increased STN excitatory drive to SNc cells result in hasten the stress-induced neurodegeneration of remaining SNc cells (fig. 14E).

**Figure 14:**
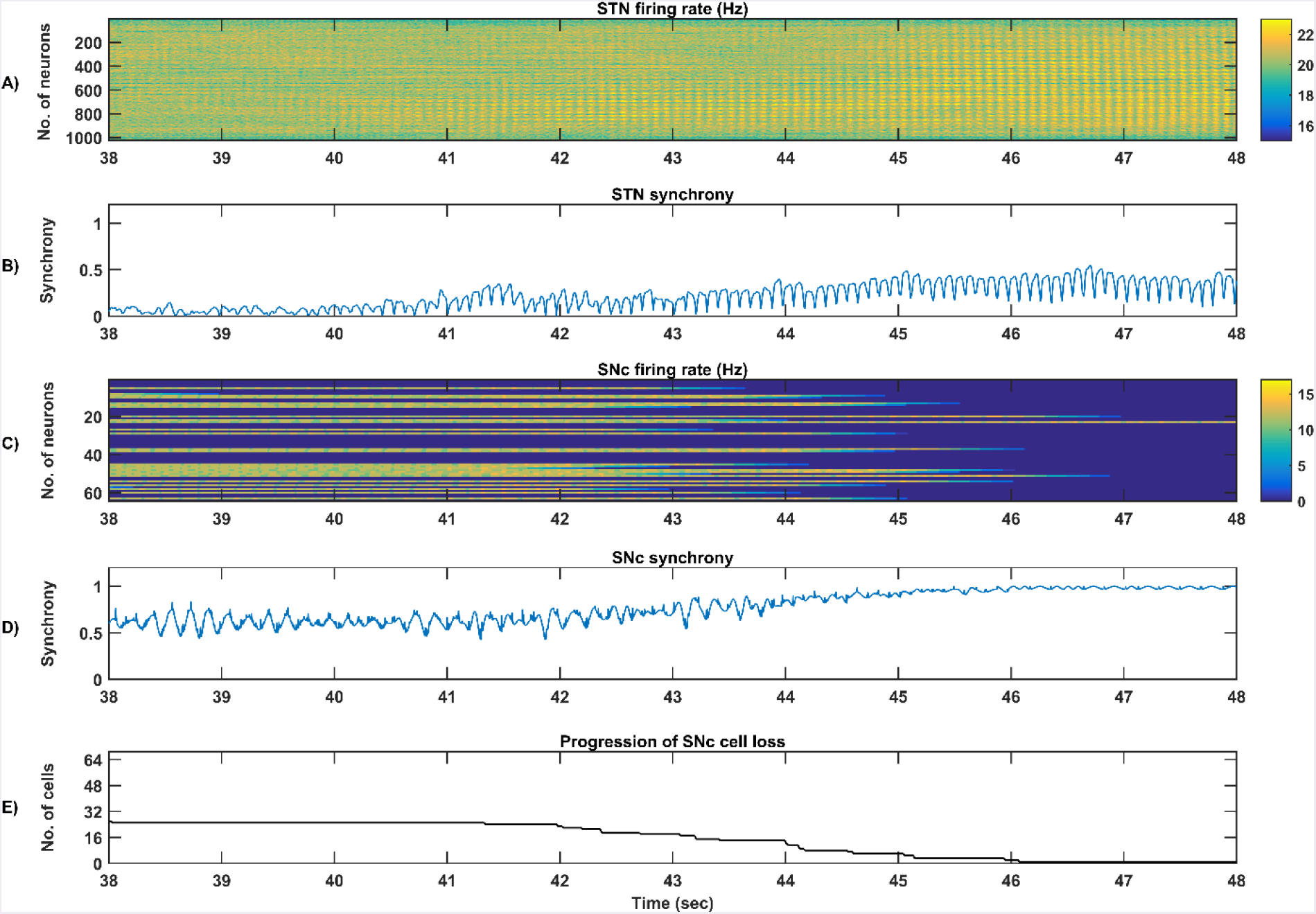
Simulation plots of STN-mediated runaway effect of neurodegeneration in SNc (Part-III in fig. 11) – Mean firing rate (1s) of STN (A) & SNc (C), Synchrony of STN (B) & SNc (D), Progression of SNc cell loss (E).

### Sensitivity of excitotoxicity model towards parameter uncertainty

To check the sensitivity of excitotoxicity model for different parametric values, we have considered two factors which can maximally influence the output results. Firstly, stress threshold (*S_thres_*) which is analogous to the apoptotic threshold and is assumed to be dependent on the amount of available energy to the cell (Albin and Greenamyre, 1992; Greene and Greenamyre, 1996). Secondly, the synaptic weight between STN and SNc (*W*_*stn*→*SNc*_) which is analogous to synaptic modification and is assumed to be modulated by the excitatory drive from STN to SNc (Hasselmo, 1994, 1997).

#### Stress threshold (*S_thres_*)

Simulation results showed that the time taken for *50%* SNc cell loss (*t*_1/2_) increases as the stress threshold increases (fig. 15A). Rate of degeneration or degeneration constant (λ) is the ratio of the number of SNc cells that degenerate in a given period of time compared with the total number of SNc cells present at the beginning of that period. The rate of degeneration (λ) decreases as the stress threshold increases as shown in the fig. 15B. These results show the importance of stress threshold in regulating excitotoxic damage to SNc and also support the idea of “weak excitotoxicity hypothesis” where SNc cells showed increased susceptibility to glutamate due to impaired cellular energy metabolism (Albin and Greenamyre, 1992; Greene and Greenamyre, 1996).

**Figure 15:**
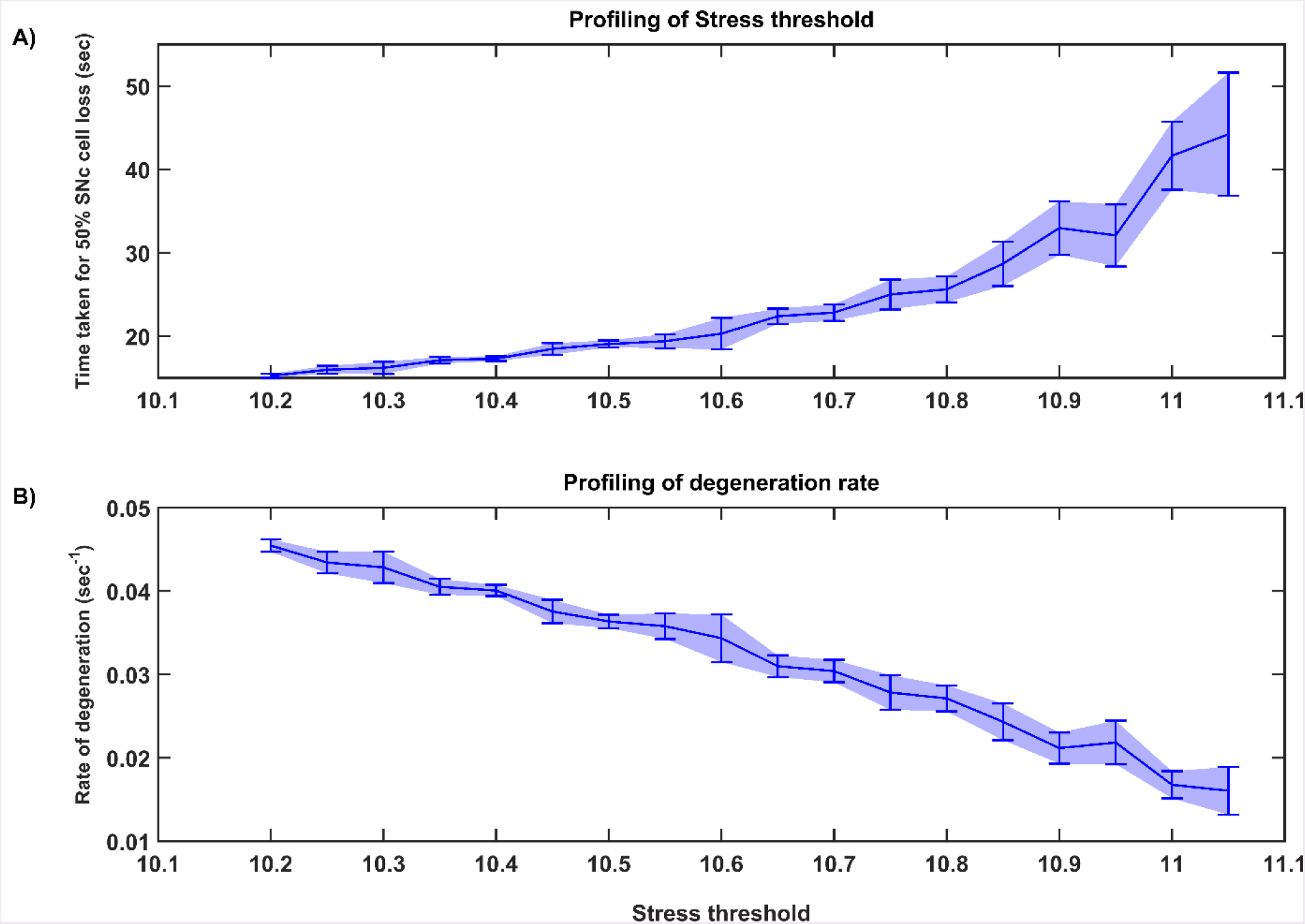
Time taken for 50% SNc cell loss for varying stress threshold (A) and Rate of degeneration with respect to the range of stress threshold (B).

#### STN-SNc synaptic weight (*W*_*STN*→*SNc*_)

Simulation results showed that time taken for *50%* SNc cell loss (*t*_1/2_) decreases as the STN-SNc synaptic weight increases as shown in the fig. 16A. The rate of degeneration (λ) increases as the STN-SNc synaptic weight increases as shown in the fig. 16B. These results show the extent of STN influence in the causation of excitotoxicity in SNc. They also support the notion that STN-mediated excitotoxicity might plays a major role in SNc cell loss in PD condition (Rodriguez et al., 1998; Blandini, 2001, 2010; Ambrosi et al., 2014).

**Figure 16:**
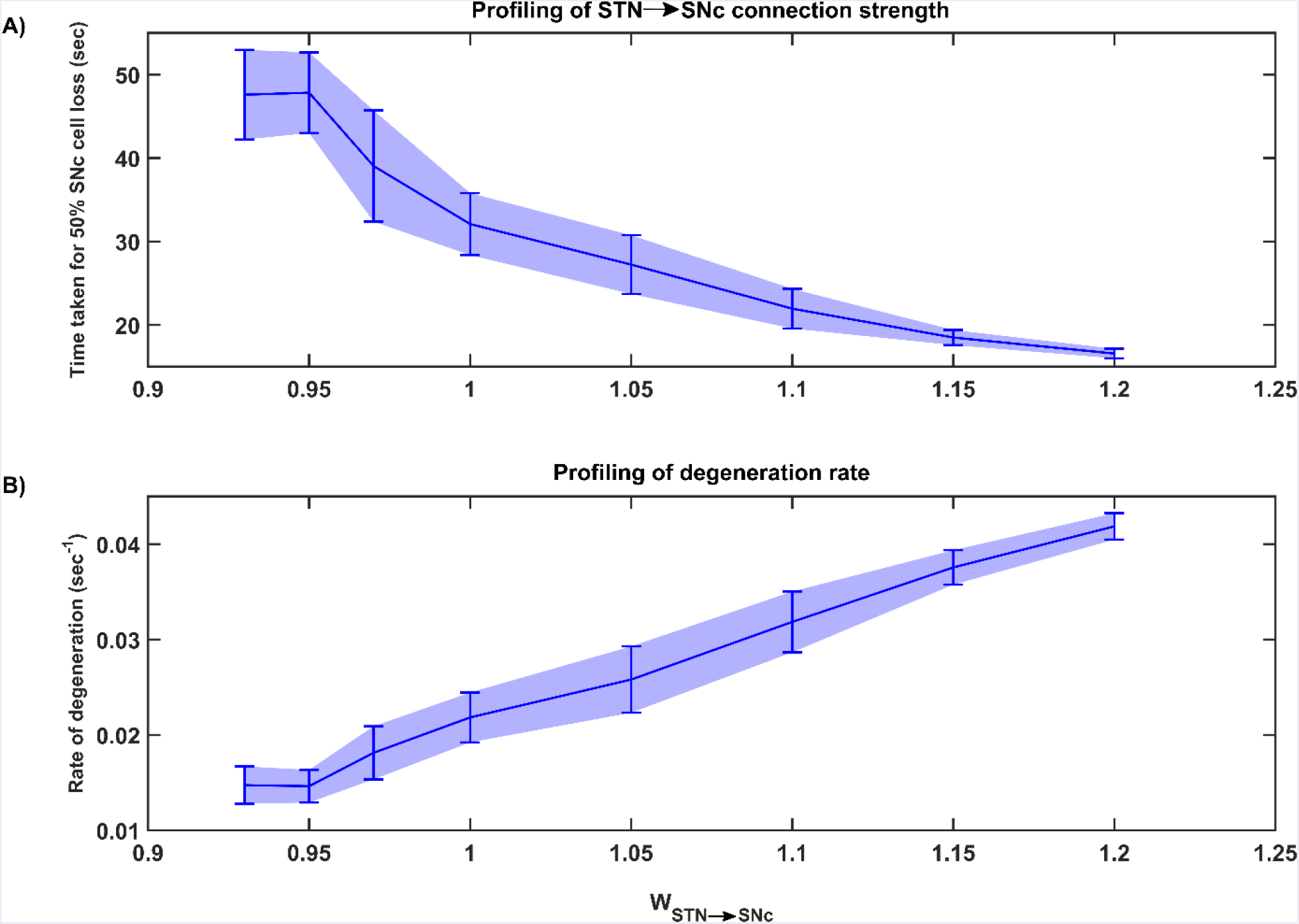
Time taken for 50% SNc cell loss for varying connection strength from STN →SNc (A) and Rate of degeneration with respect to the range of connection strength (B).

### Strategies for neuroprotection of SNc

We now extend the proposed excitotoxic model to study the effect of various therapeutic interventions on SNc cell loss. The following three types of interventions are simulated: 1) drugs, 2) surgical interventions, and 3) Deep Brain Stimulation (DBS).

#### Glutamate inhibition therapy

The effect of glutamate agonists and antagonists on the progression of SNc cell loss was implemented in as the manner specified in the methods section. The onset of glutamate therapy at different stages of SNc cell loss showed that cell loss was delayed or halted as shown in the fig. 17-19. For the glutamate therapy which is initiated at 25%, 50%, and 75% SNc cell loss, the progression of SNc cell loss was halted when the percentage of glutamate inhibition administrated was above 50%. As the glutamate dosage increases the progression of SNc cell loss delays and after a certain dosage of glutamate inhibitors the SNc cell loss halts. There was no change in the course of SNc cell loss for low levels of glutamate inhibition (fig. 17A(iii)). The peak of the instantaneous rate of degeneration decreases as the therapeutic intervention delayed in the case of 10% glutamate inhibition.

**Figure 17:**
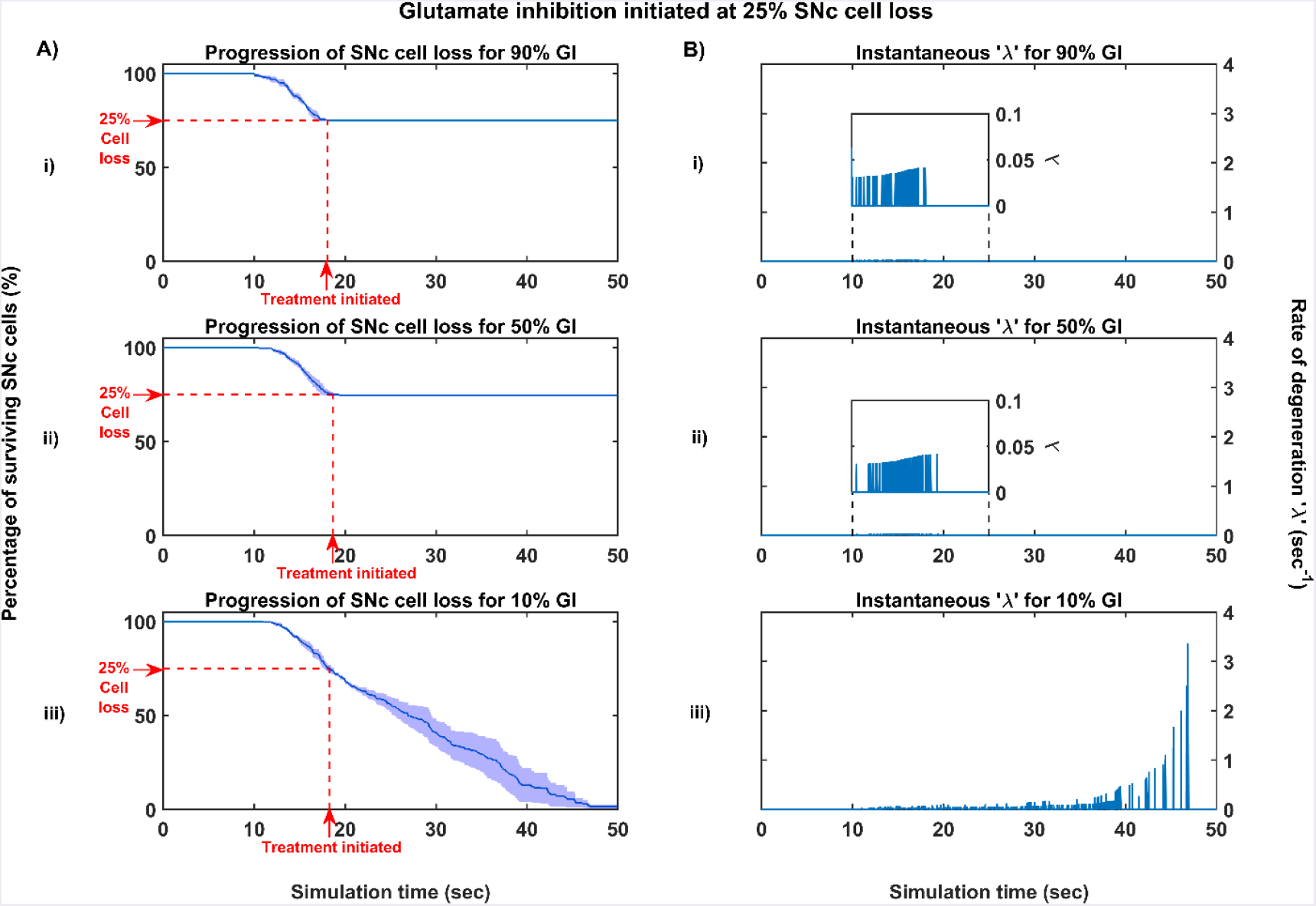
Simulation plots for glutamate inhibition (GI) initiated at 25% SNc cell loss. (A) Progression of SNc cell loss for (i) 90% (ii) 50% and (iii) 10% GI, (B) Instantaneous rate of degeneration for (i) 90% (ii) 50% and (iii) 10% GI.

**Figure 18:**
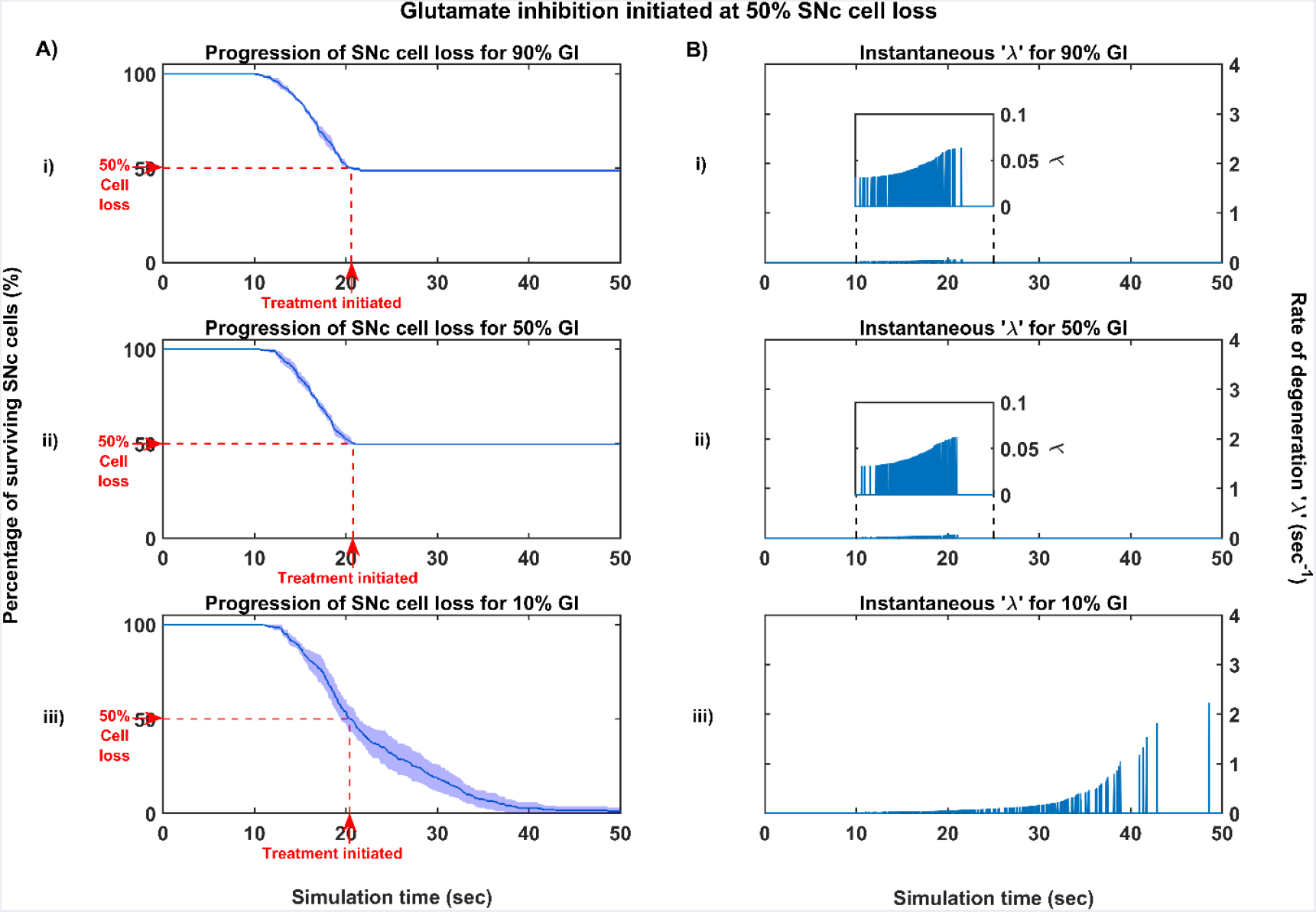
Simulation plots for glutamate inhibition (GI) initiated at 50% SNc cell loss. (A) Progression of SNc cell loss for (i) 90% (ii) 50% and (iii) 10% GI, (B) Instantaneous rate of degeneration for (i) 90% (ii) 50% and (iii) 10% GI.

**Figure 19:**
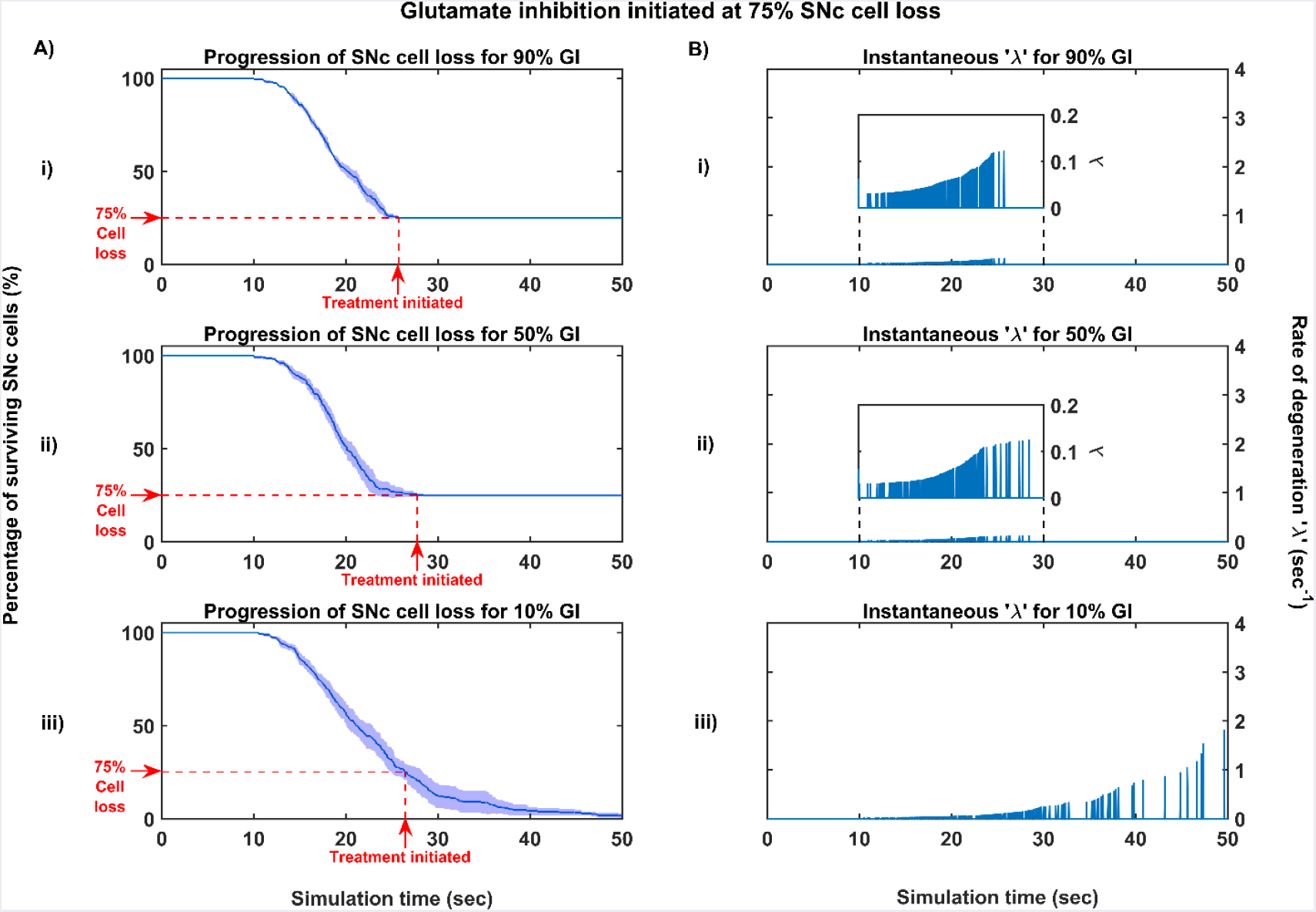
Simulation plots for glutamate inhibition (GI) initiated at 75% SNc cell loss. (A) Progression of SNc cell loss for (i) 90% (ii) 50% and (iii) 10% GI, (B) Instantaneous rate of degeneration for (i) 90% (ii) 50% and (iii) 10% GI.

#### Dopamine restoration therapy

The effect of dopamine agonists on the progression of SNc cell loss was also implemented in the manner specified in the methods section. The onset of dopamine agonist therapy at different stages of SNc cell loss showed that progression of cell loss was delayed or halted as shown in the fig. 20-22. For the dopamine agonists therapy which is initiated at 25%, 50%, and 75% SNc cell loss, the progression of SNc cell loss was delayed when the percentage of dopamine restoration was mere 10%. The neuroprotective effect of dopamine agonist therapy is dependent on the level of restoration of dopamine tone on the STN. In other words, as the dopamine content in STN increases, the progression of SNc cell loss delays and halts only when dopamine content is almost completely restored. Unlike glutamate inhibition, the progression of SNc cell loss wasn’t halted even at 100% dopamine restored in case of intervention at 25% and 50% cell loss. But in the case of intervention at 75% cell loss, the progression of SNc cell loss halted for 100% and 50% but not for 10% dopamine restoration.

**Figure 20:**
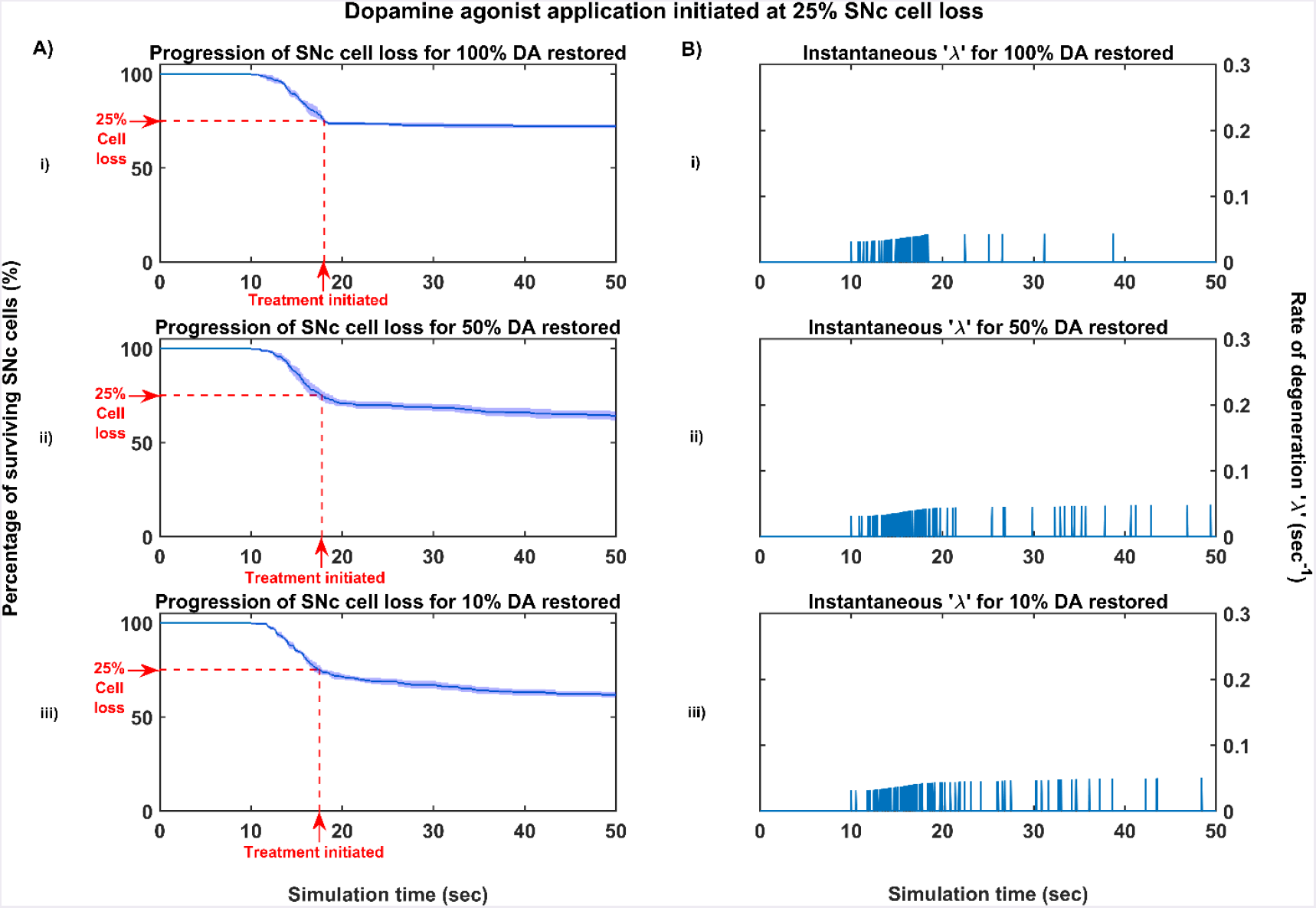
Simulation plots for dopamine agonist application (DAA) initiated at 25% SNc cell loss. (A) Progression of SNc cell loss for (i) 100% (ii) 50% and (iii) 10% DAA, (B) Instantaneous rate of degeneration for (i) 100% (ii) 50% and (iii) 10% DAA.

**Figure 21:**
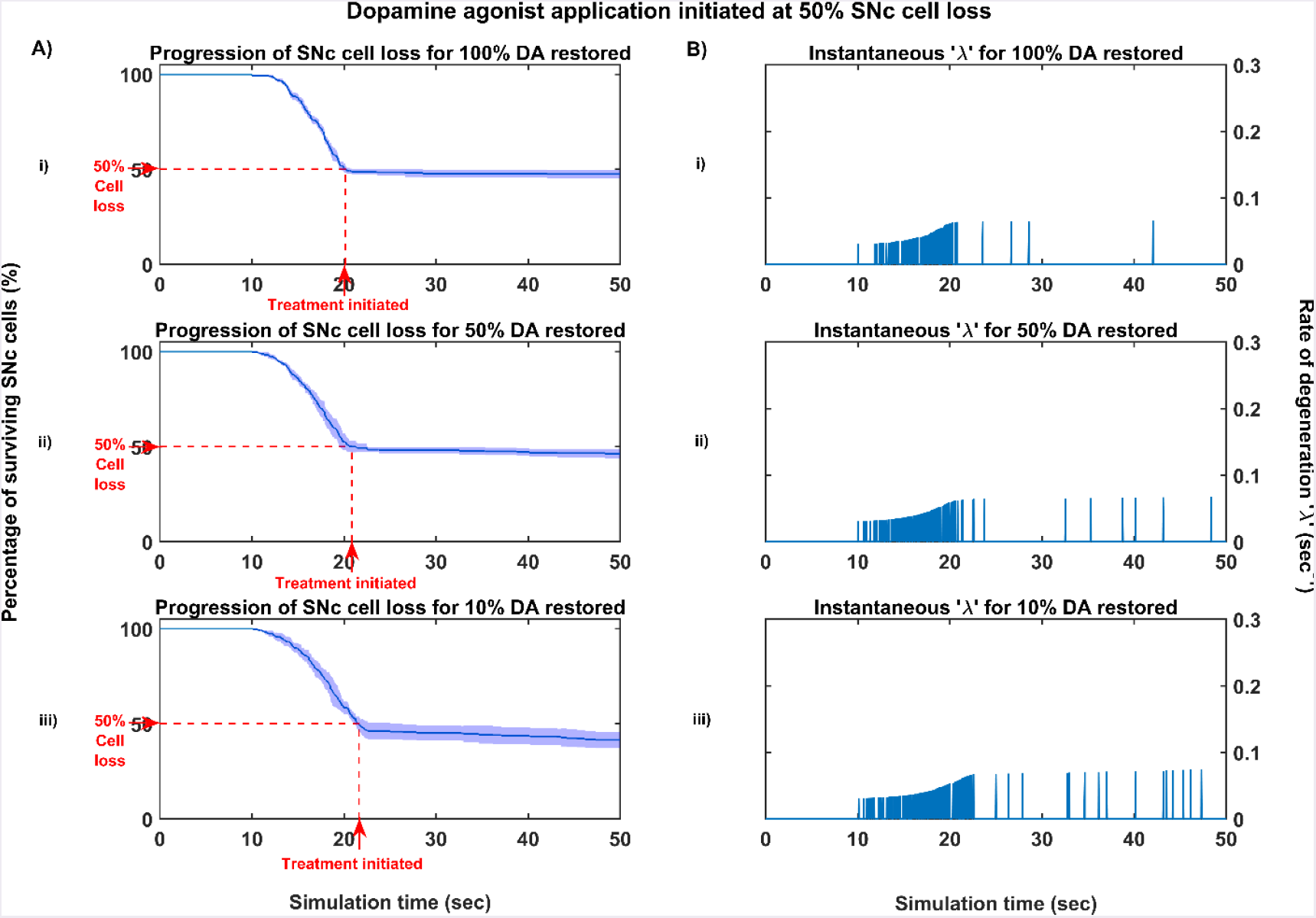
Simulation plots for dopamine agonist application (DAA) initiated at 50% SNc cell loss. (A) Progression of SNc cell loss for (i) 100% (ii) 50% and (iii) 10% DAA, (B) Instantaneous rate of degeneration for (i) 100% (ii) 50% and (iii) 10% DAA.

**Figure 22:**
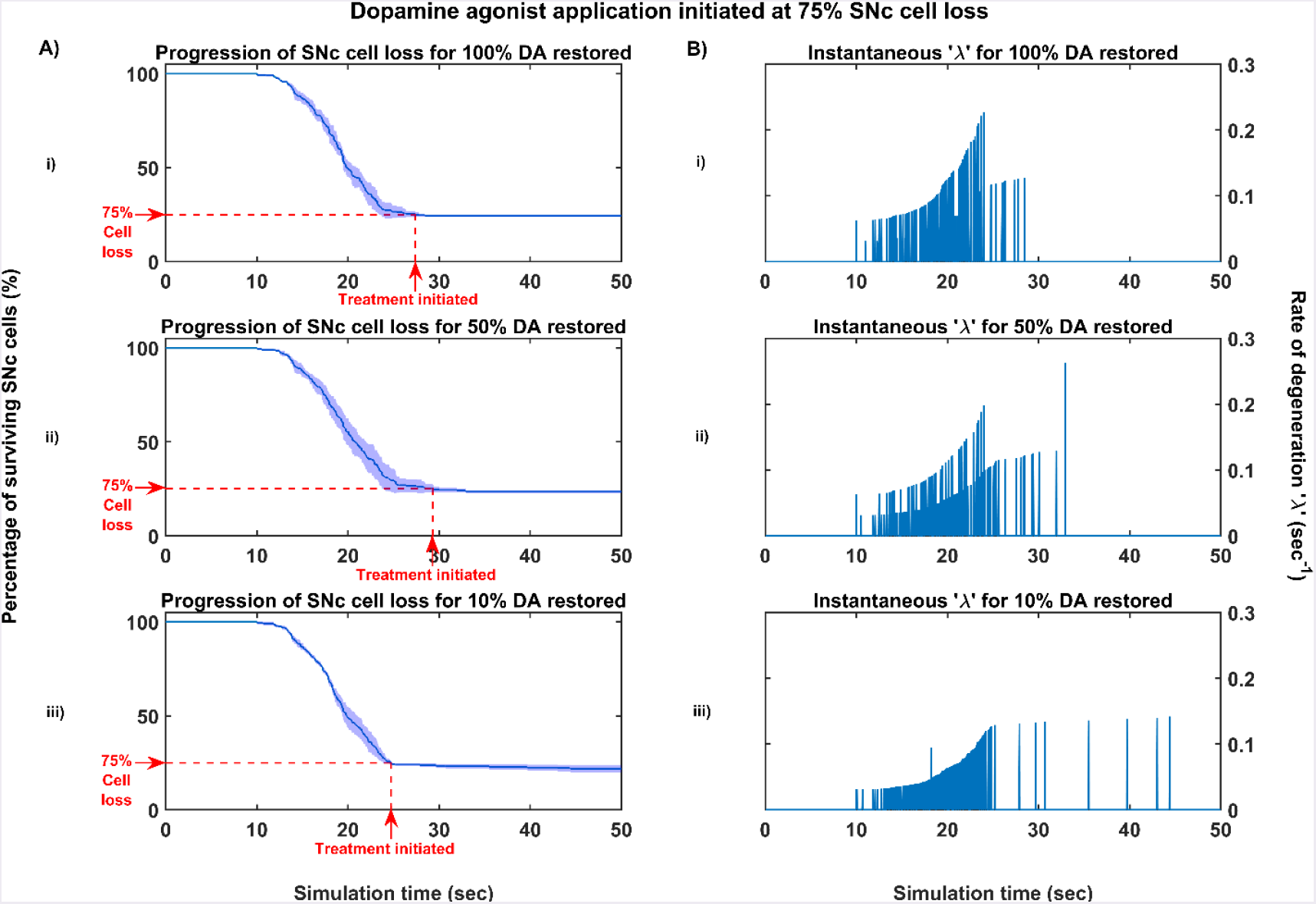
Simulation plots for dopamine agonist application (DAA) initiated at 75% SNc cell loss. (A) Progression of SNc cell loss for (i) 100% (ii) 50% and (iii) 10% DAA, (B) Instantaneous rate of degeneration for (i) 100% (ii) 50% and (iii) 10% DAA.

#### Subthalamotomy

The effect of subthalamotomy on the progression of SNc cell loss was implemented in a way described in the methods section. The onset of STN ablation therapy at different stages of SNc cell loss showed that progression of cell loss was delayed or halted (fig. 23-25). The neuroprotective effect of subthalamotomy is dependent on the proportion of lesioning of STN population. In other words, as the proportion of STN lesioning increases the progression of SNc cell loss delays and halts only when almost all of the STN population is lesion. The progression of SNc cell loss was halted only at 100% STN lesioning in all cases of intervention. But as the proportion of STN lesioning decreases, rate of degeneration increases as shown in the figs. 23B, 24B, 25B.

**Figure 23:**
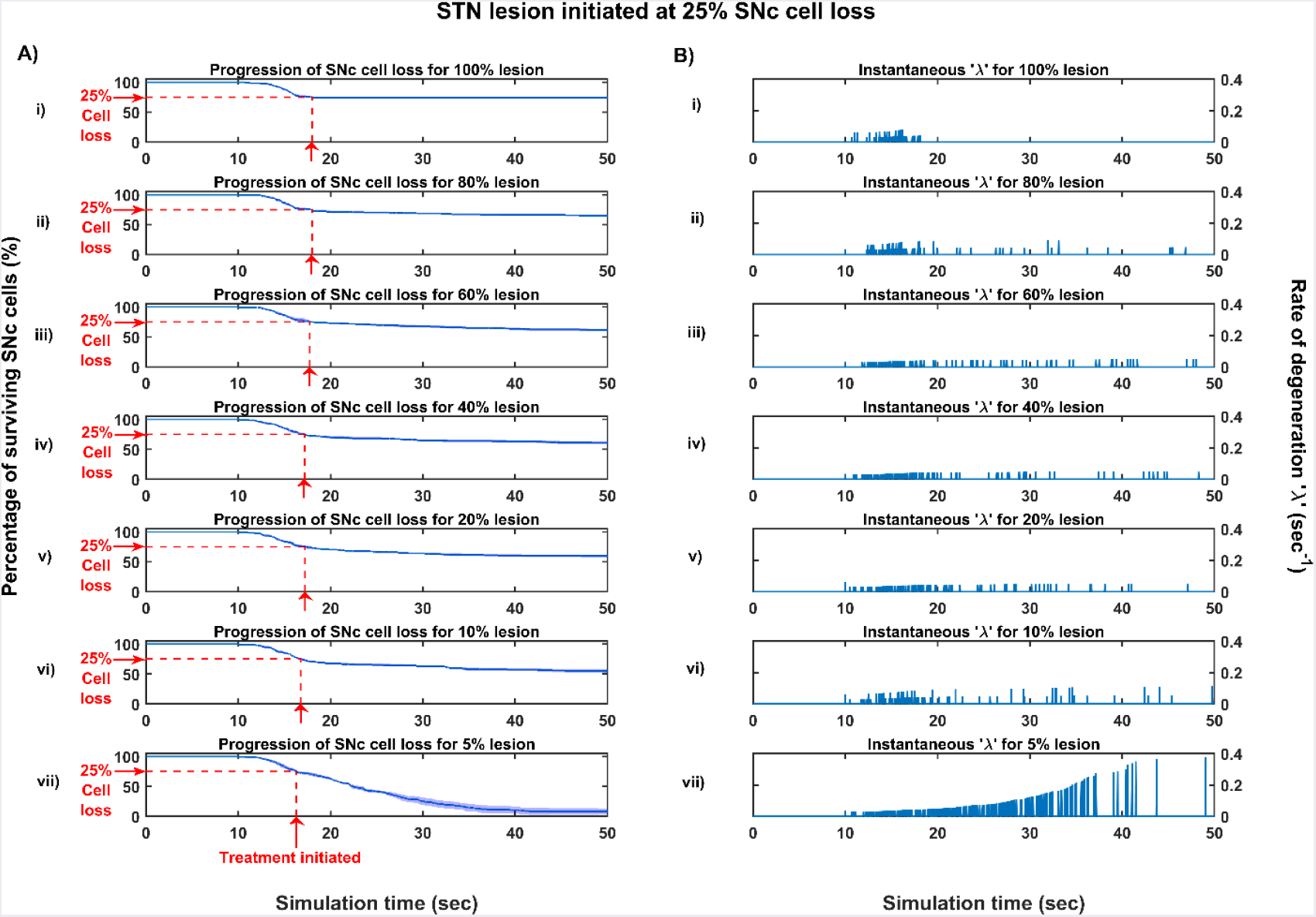
Simulation plots for STN lesion (LES) initiated at 25% SNc cell loss. (A) Progression of SNc cell loss for (i) 100% (ii) 80% (iii) 60% (iv) 40% (v) 20% (vi) 10% and (vii) 5% LES, (B) Instantaneous rate of degeneration for (i) 100% (ii) 80% (iii) 60% (iv) 40% (v) 20% (vi) 10% and (vii) 5% LES.

**Figure 24:**
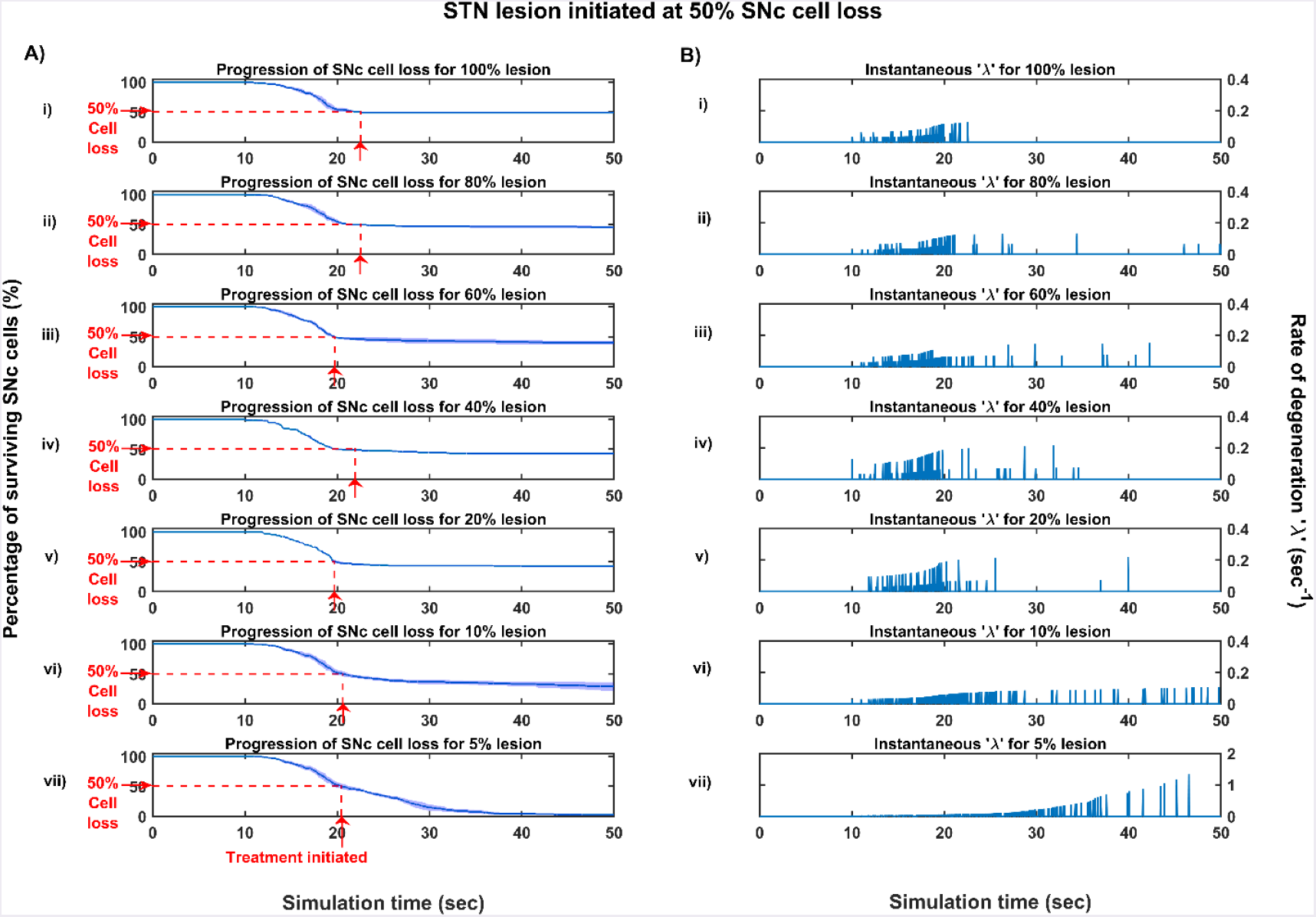
Simulation plots for STN lesion (LES) initiated at 50% SNc cell loss. (A) Progression of SNc cell loss for (i) 100% (ii) 80% (iii) 60% (iv) 40% (v) 20% (vi) 10% and (vii) 5% LES, (B) Instantaneous rate of degeneration for (i) 100% (ii) 80% (iii) 60% (iv) 40% (v) 20% (vi) 10% and (vii) 5% LES.

**Figure 25:**
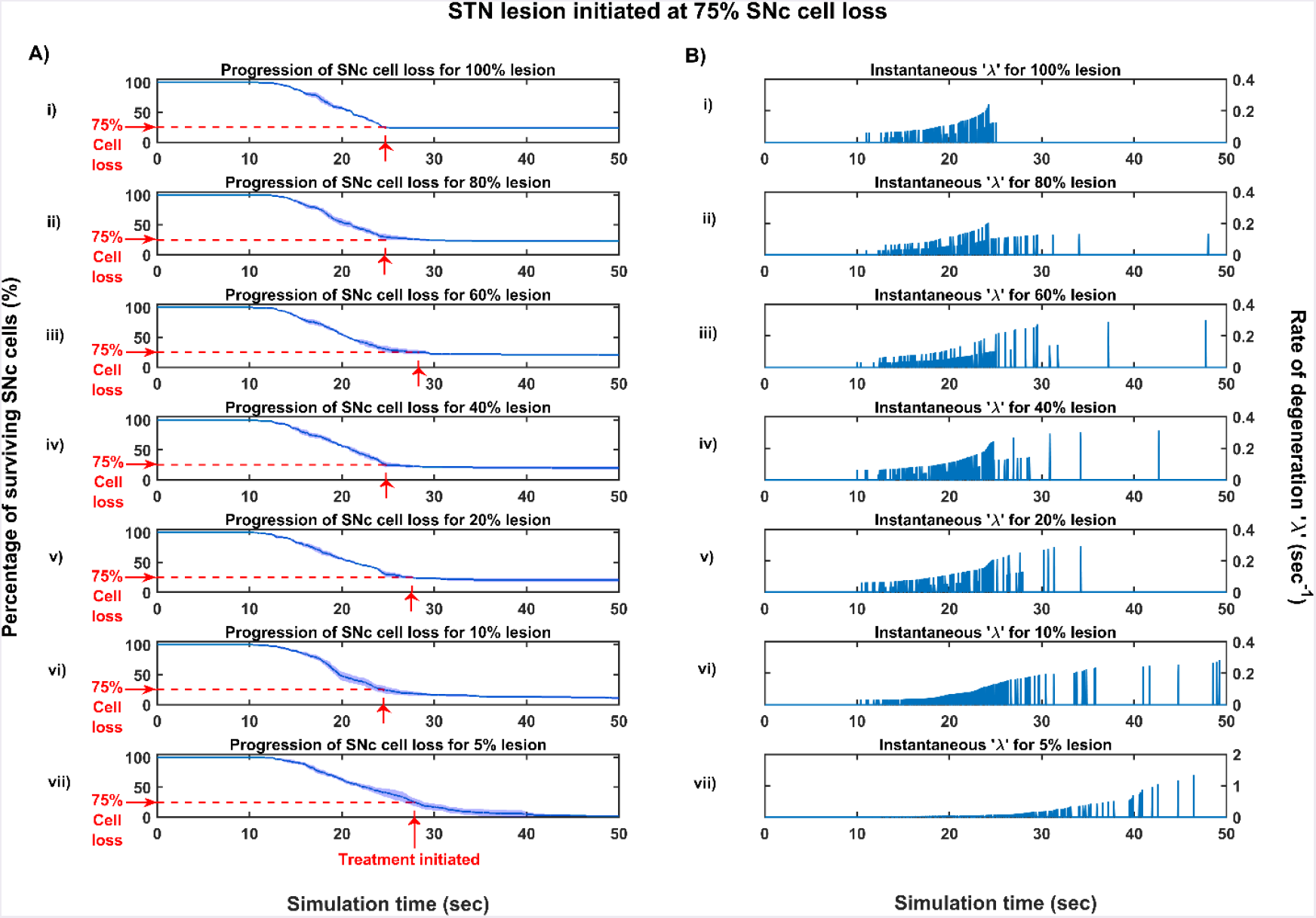
Simulation plots for STN lesion (LES) initiated at 75% SNc cell loss. (A) Progression of SNc cell loss for (i) 100% (ii) 80% (iii) 60% (iv) 40% (v) 20% (vi) 10% and (vii) 5% LES, (B) Instantaneous rate of degeneration for (i) 100% (ii) 80% (iii) 60% (iv) 40% (v) 20% (vi) 10% and (vii) 5% LES.

#### Deep brain stimulation of STN

The effect of deep brain stimulation on the progression of SNc cell loss was implemented in the way described in the methods section. Along with the stimulation of STN, the inhibitory drive to STN through the afferent connections as result of antidromic activation of the GPe population and the synaptic depression in STN as result of increased axonal and synaptic failures in STN were incorporated in the model.

As specified earlier, different stimulation configurations and stimulus waveforms were implemented while exploring the optimal DBS parameters for therapeutic benefits. The STN population response for different types of DBS protocol was shown in the figs. 26-28. To study the neuroprotective effect, stimulation parameters which reduce the STN overactivity (Meissner et al., 2005) due to dopamine depletion were chosen. The biphasic stimulus pulse shows more therapeutic benefits than monophasic stimulus pulse; biphasic current alleviates the STN pathological activity without increasing the firing rate of STN population as a whole. The four-contact point type of stimulation configuration required lesser stimulus amplitude for producing the same effect when compared with the other two configurations. From these preliminary studies, we can say that four-contact point configuration with biphasic stimulus pulse gives maximum therapeutic benefits from the neuroprotective point of view. For DBS parameters, see the *Table-2*.

**Figure 26:**
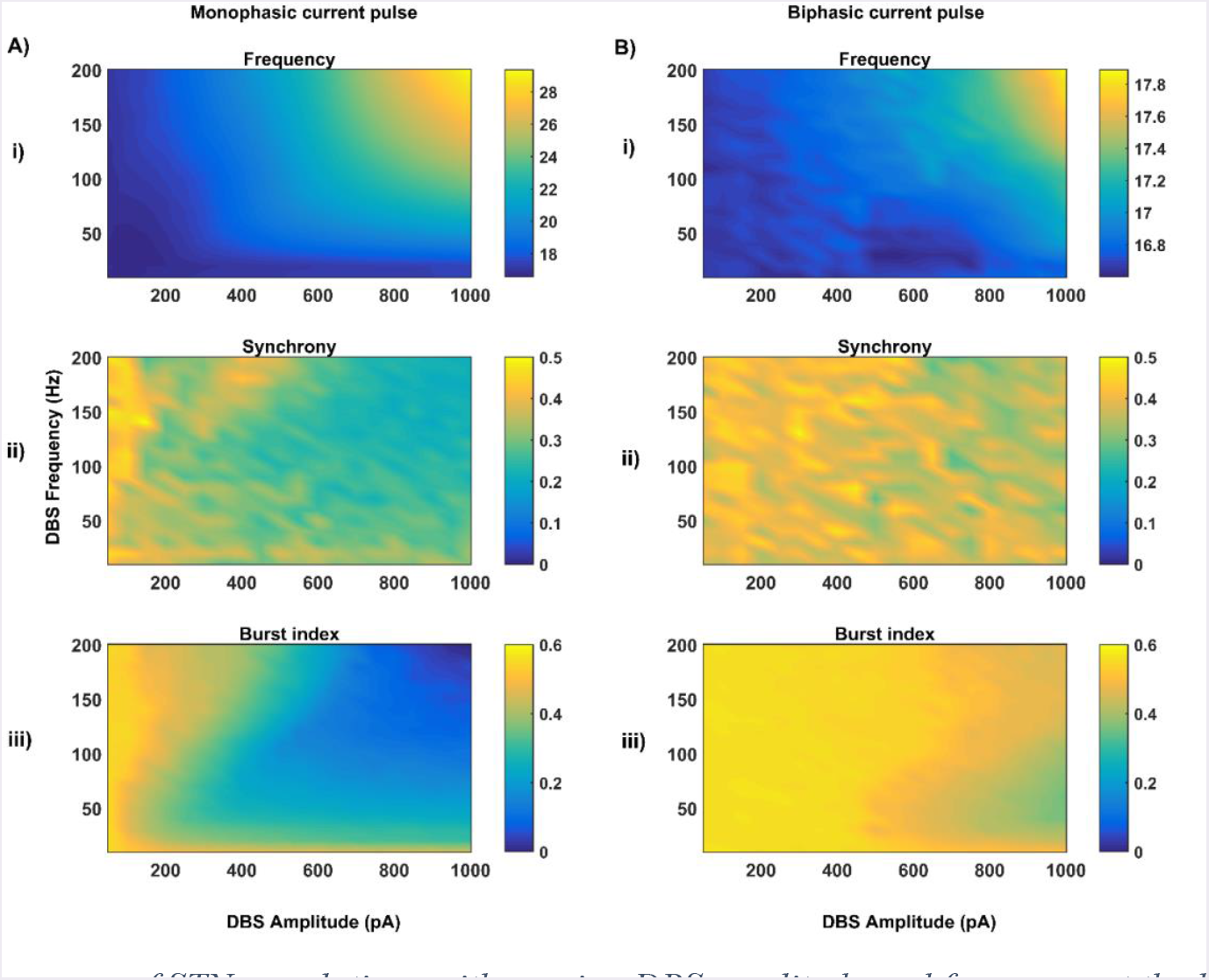
The response of STN populations with varying DBS amplitude and frequency at the level of network properties (Frequency (i), Synchrony (ii) and Burst index (iii)) for single contact point (SCP) monophasic (A) and Biphasic (B) current pulses.

**Figure 27:**
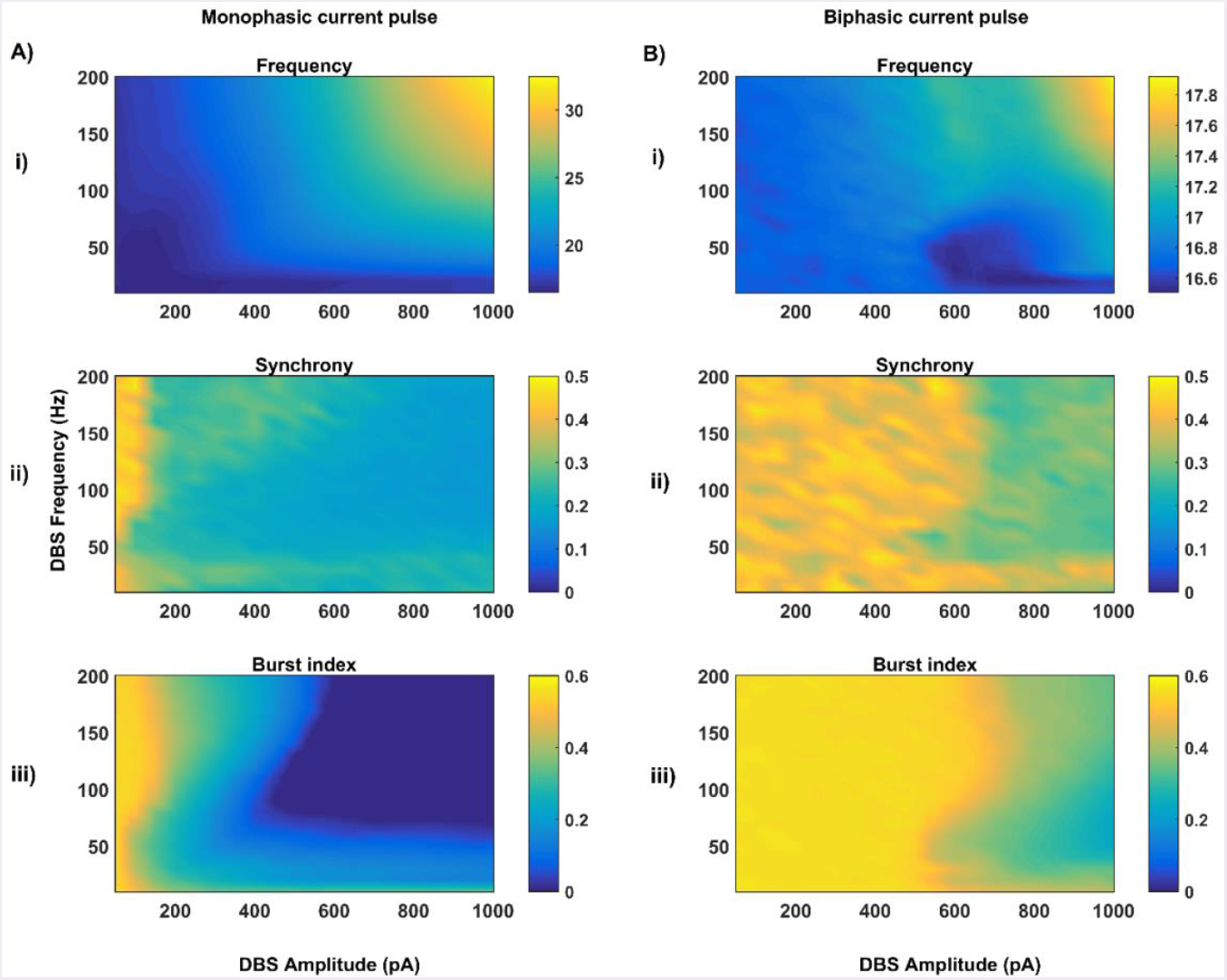
The response of STN populations with varying DBS amplitude and frequency at the level of network properties (Frequency (i), Synchrony (ii) and Burst index (iii)) for four contact point (FCP) monophasic (A) and Biphasic (B) current pulses.

**Figure 28:**
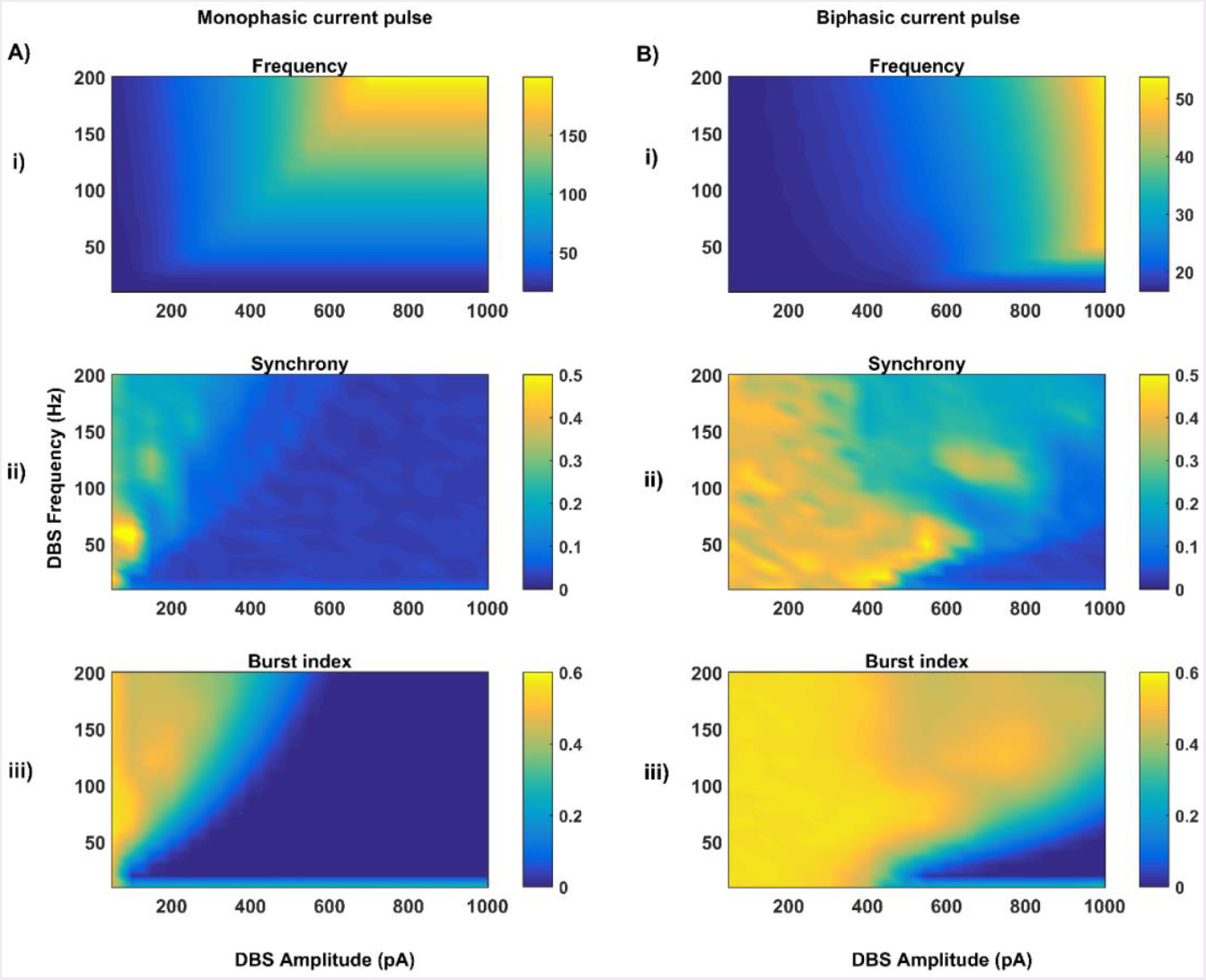
The response of STN populations with varying DBS amplitude and frequency at the level of network properties (Frequency (i), Synchrony (ii) and Burst index (iii)) for multiple contact point (MCP) monophasic (A) and Biphasic (B) current pulses.

**Table-2.** DBS parameter values obtained from the preliminary studies

| Parameter(s) | SCP | FCP | MCP |
| --- | --- | --- | --- |
| DBS frequency ($f_{DBS}$) in Hertz | 130 | 130 | 130 |
| Monophasic pulse width ($\delta_{DBS}$) in microseconds | 100 | 100 | 100 |
| Biphasic pulse width ($\delta_{DBS}$) in microseconds | 200 | 200 | 200 |
| Monophasic DBS amplitude ($A_{DBS}$) in picoampere | 650 | 650 | 650 |
| Biphasic DBS amplitude ($A_{DBS}$) in picoampere | 1000 | 1000 | 1000 |
| Spread of the current ($\sigma_{DBS}$) | 5 | 2 | 0 |
| Electrode contact point(s) | (16,16) | (8,8)<br>(8,24)<br>(24,8)<br>(24,24) | Many |

#### Neuroprotective effect of DBS

To understand the neuroprotective therapeutic mechanism of DBS in PD (Benazzouz et al., 2000; Maesawa et al., 2004; Temel et al., 2006; Wallace et al., 2007; Spieles-Engemann et al., 2010; Musacchio et al., 2017), we have investigated some of the dominant hypotheses regarding therapeutic effect of DBS viz., 1) excitation hypothesis, 2) inhibition hypothesis and most recent one 3) disruptive hypothesis (McIntyre et al., 2004; Chiken and Nambu, 2015).

The excitation hypothesis was implemented by direct stimulation of the STN population in the proposed excitotoxicity model. The simulation results show that DBS to STN diminishes the pathological synchronized activity but in turn increases the firing rate of the STN population which wasn’t apt for neuroprotection.

Next, we have implemented the inhibition hypothesis where antidromic activation of GPe neurons during DBS is highlighted, thereby increasing the inhibitory drive to STN (Mandali and Chakravarthy, 2016). In this scenario also, the inhibitory drive from GPe wasn’t sufficient to produce comprehensive neuroprotection in as shown in the fig. 29, 30. On average FCP stimulus configuration produced better neuroprotective effect compared to other two configurations in both monophasic and biphasic current (fig. 29B, 30B). And MCP stimulus configuration results in worsening the disease progression by hasten the SNc cell loss in monophasic stimulus (fig. 29C) but in biphasic stimulus neuroprotection increased with higher levels of antidromic activation in all stages of therapeutic intervention (fig. 30C).

**Figure 29:**
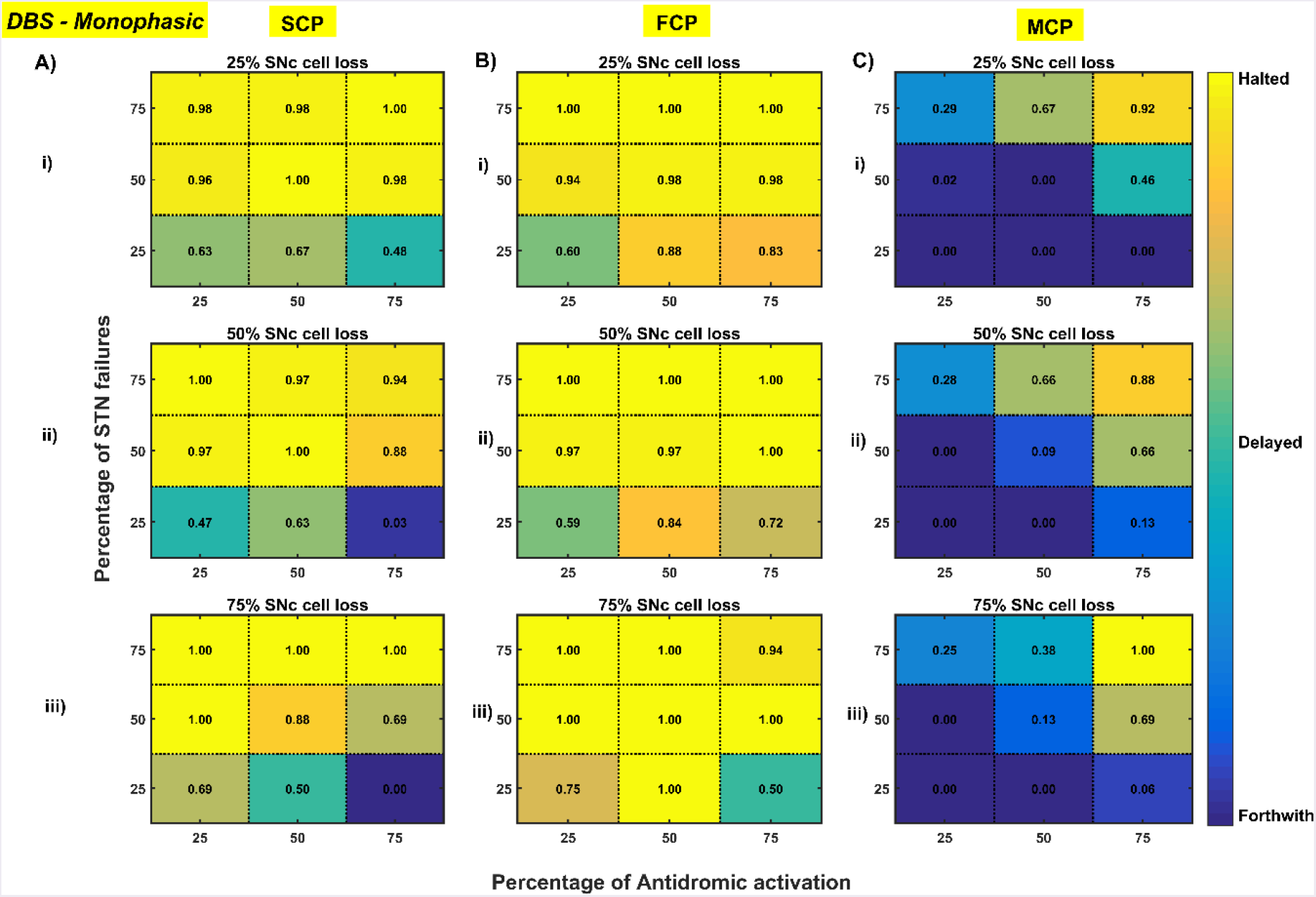
Profiling of monophasic stimulus waveform for different stimulation configuration in order to achieve the maximal neuroprotective effect of DBS.

**Figure 30:**
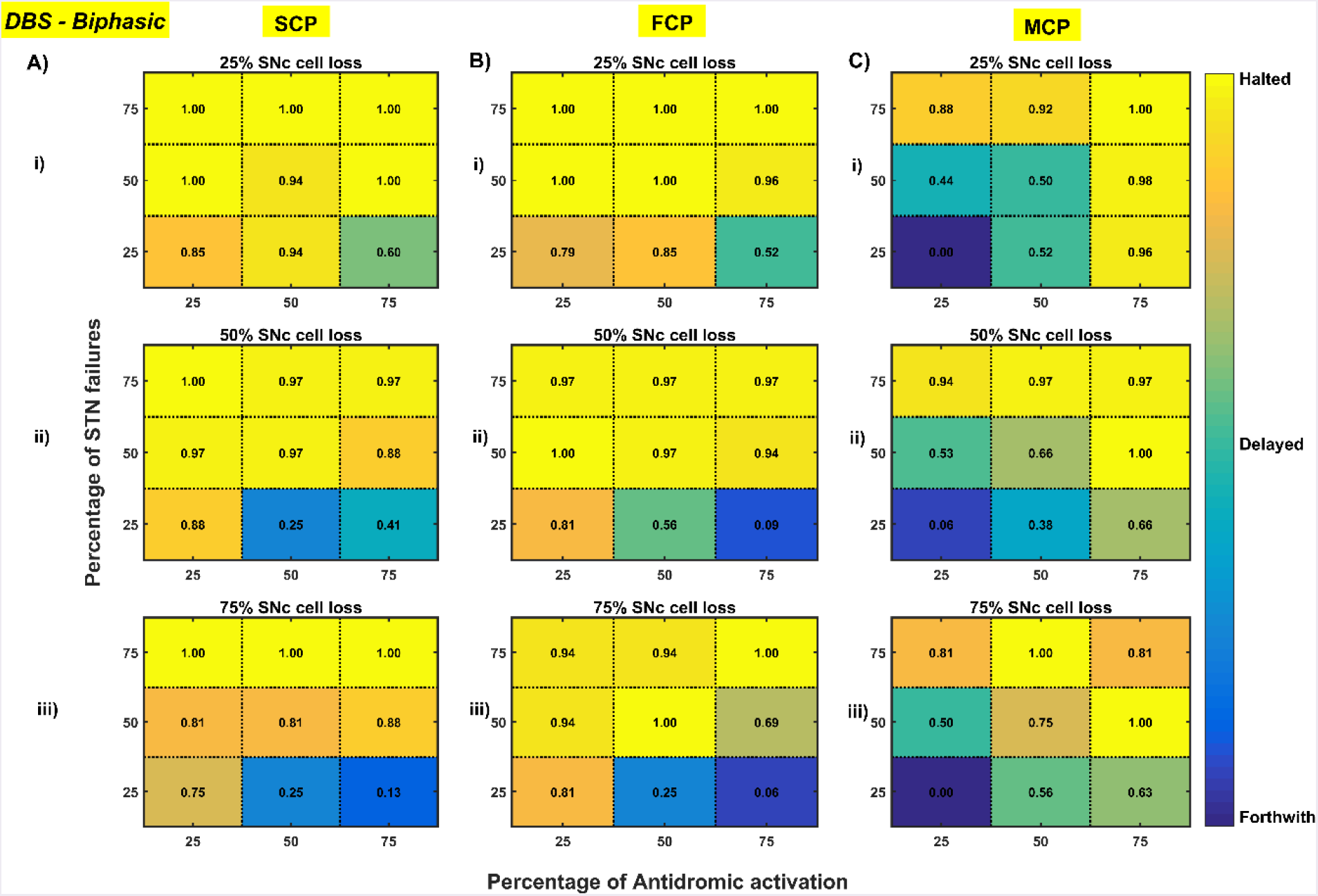
Profiling of biphasic stimulus waveform for different stimulation configuration in order to achieve the maximal neuroprotective effect of DBS.

Finally, the disruptive hypothesis was implemented by increasing the proportion of axonal and synaptic failures in STN population (Rosenbaum et al., 2014). From simulation results, it was observed that the progression of SNc cell loss was delayed or halted as the percentage of STN axonal and synaptic failures increased as shown in the fig. 29, 30. On average FCP stimulus configuration produced better neuroprotective effect compared to other two configurations in both monophasic and biphasic current (fig. 29B, 30B). For the even higher percentage of STN axonal and synaptic failures, the neuroprotective effect was not pronounced in monophasic MCP DBS setting but in biphasic MCP DBS setting neuroprotection increased with the higher percentage of STN axonal and synaptic failures (fig. 30C).

## IV. Discussion & Conclusions

### Excitotoxicity model

The goal of this work was to develop a model which investigates the role of excitotoxicity in SNc cell loss, where excitotoxicity was caused by STN and precipitated by energy deficiency. The study suggests that excitotoxicity in SNc is initially driven by an energy deficit which leads to initial dopamine reduction as a result of SNc cell loss. This initial dopamine reduction causes disinhibition of STN which in turns leads to excitotoxic damage to its target nuclei including SNc (Rodriguez et al., 1998). The excitotoxicity which was driven by energy impairment, termed as “weak excitotoxicity,” results in increased vulnerability of SNc neurons to even physiological concentration of glutamate. The excitotoxicity which was driven by overactive excitatory STN neurons termed as “strong excitotoxicity” results in overactivation of glutamatergic receptors on SNc neurons (Albin and Greenamyre, 1992; Rodriguez et al., 1998). In summary, it appears that the excitotoxic cause of SNc cell loss in PD might be initiated by weak excitotoxicity mediated by energy deficit, and followed by strong excitotoxicity, mediated by disinhibited STN.

It was observed that SNc neurons are selectively vulnerable in PD as a result of their characteristics like unique massive unmyelinated axonal arbors, at least two-fold more synapses compared to other basal ganglia nuclei, the presence of reactive neurotransmitter namely dopamine, calcium loading, pacemaking, higher basal metabolic rate etc. (Mosharov et al., 2009; Bolam and Pissadaki, 2012; Pissadaki and Bolam, 2013; Dragicevic et al., 2015b; Pacelli et al., 2015). These features of SNc neurons place tight constraints on the maintenance of metabolic homeostasis. Any imbalance in energy supply and demand results in energy deficits in SNc, a development that perhaps can eventually lead to neurodegeneration (Wellstead and Cloutier, 2011; Pissadaki and Bolam, 2013).

The results from the proposed model reinforces the role of STN in regulating SNc cell loss (Hamani et al. 2004, 2017; Hammond et al. 1978; Iribe et al. 1999; Hitoshi Kita and Kitai 1987; Meissner et al. 2003; Mintz et al. 1986; Paquet et al. 1997; Smith and Grace 1992; Smith, Charara, and Parent 1996). The model results show that although cell loss was observed, there was no increased synchrony in the STN population as shown in fig. 13B which is a pathological marker of the PD condition (Lintas et al., 2012; Yang et al., 2016). Thus, the SNc cell loss and STN synchrony have a threshold-like relation where there is an increased STN synchrony only after substantial SNc cell loss. The initial SNc cell loss leads to further activation of STN by disinhibition, which in turn further activates SNc compensating for the dopamine loss, acting as a pre-symptomatic compensatory mechanism (Bezard et al., 2003). From experimental literature, it was reported that onset of PD symptoms occurs only after there is more than 50% SNc cell loss (Bezard et al., 2001; Lang and Obeso, 2004; Toulouse and Sullivan, 2008). This was observed in our simulation results also where around 50-70% SNc cell loss results in an increased STN synchrony. As a result of substantial SNc cell loss, decreased dopamine causes disinhibition of STN which in turn overactivates STN, eventually producing a runaway effect (Hasselmo, 1994, 1997) wherein SNc cells are lost continually due to excitotoxic damage (Rodriguez et al., 1998). The threshold-like behavior of SNc cell loss and STN synchrony might be facilitated by the inhibitory drive from GPe to STN as result of the proliferation of GPe-STN synapses (Fan et al., 2012) which also acts as a presymptomatic compensatory mechanism. It was also reported that lesioning of GPe caused progressive SNc cell loss by increasing STN activity (Wright et al., 2002) and lesioning of STN proven to be neuroprotective (Wright and Arbuthnott, 2007).

To summarize, up to a point of stress threshold, SNc cells can survive indefinitely; but if, for any reason, there is loss of cells in SNc, and the SNc cell count falls below a threshold, from that point onwards, the aforementioned runaway effect kicks in leading to a progressive and irrevocable cell loss. Such cell loss is strongly reminiscent of cell loss due to neurodegeneration.

### Neuroprotective therapy

#### Glutamate inhibition therapy

The glutamate inhibition therapy was successful in delaying or halting the progression of SNc cell loss by inhibiting the excitatory drive from STN to SNc, thereby diminishing STN-mediated excitotoxicity. The neuroprotective effect of glutamate inhibition therapy was dependent on the dosage of glutamate inhibitors, analogous to the trend seen in the experimental data (Austin et al., 2010). As the disease progresses, the effect of glutamate inhibition on the rate of degeneration increases. Baicalein, a Chinese medicine extracted from the root of *Scutellaria baicalensis Georgi* was able the show neuroprotective properties; this drug diminishes excitotoxicity by inhibiting the release of glutamate and antagonizing NMDA receptors (Lee et al., 2003; Li et al., 2017); it also diminishes excitotoxicity-induced by glucose-deprivation (Lee et al., 2003). Our modeling studies suggest that the selective glutamate inhibitors would show neuroprotective effect all through the course of the disease and strikingly even in advanced stages of disease progression.

#### Dopamine restoration therapy

The dopamine restoration therapy was successful in delaying or halting the progression of SNc cell loss by restoring the dopamine tone to the dopamine-deprived brain which in turn restores inhibition of STN, thereby diminishing STN-mediated excitotoxicity. The neuroprotective effect of dopamine restoration therapy was dependent on the extent of dopamine restored in modulating dopamine tone on the STN. Olanow and co-workers propose that the neuroprotective effect of dopamine agonists can be accounted by any of the following mechanisms: LDOPA sparing, activation of autoreceptors, antioxidant effects, antiapoptotic effects and finally by ameliorating STN-mediated excitotoxicity (Olanow et al., 1998; Schapira and Olanow, 2003; Piccini and Pavese, 2006). From our computational study, the neuroprotective effect of dopamine agonists appears to be due to the amelioration of STN-mediated excitotoxicity by restoring the dopamine tone to STN population. Recently, Vaarmann and co-workers conducted an experiment on midbrain cell cultures where they showed the neuroprotective effect of dopamine agonists against glutamate-induced excitotoxicity (Vaarmann et al., 2013). Our modeling studies suggest that the dopamine agonist therapy would show neuroprotective effect not only in early stages but also in the late stages of disease progression with reduced effect. In the late stages of disease progression, the neuroprotective effect of glutamate inhibition therapy is more prominent than dopamine restoration therapy from our computational study.

#### Subthalamotomy

Subthalamotomy was successful in delaying or halting the progression of SNc cell loss which is similar to glutamate inhibition therapy by reducing the excitatory drive from STN to SNc thereby diminishing STN-mediated excitotoxicity. The neuroprotective effect of subthalamotomy was dependent on the proportion of STN lesioned. From our study, it can be said that STN ablation mostly delays the progression of SNc cell loss but very rarely halts it. This phenomenon was not much evident in the late stages of disease progression which is consistent with the standard clinical understanding that neuroprotection therapies in the late-stages of PD are not fruitful (Olanow et al., 2009; Guridi and Obeso, 2015). Early treatment with subthalamotomy in PD can have a neuroprotective effect (Jourdain et al., 2014; Guridi and Obeso, 2015; Guridi et al., 2016) a trend that is reflected in our computational study.

Another factor underlying the neuroprotective effect of subthalamotomy during the early stage of PD is the involvement of presymptomatic compensation mechanisms (Bezard et al., 2003; Blesa et al., 2017). One of the compensatory mechanisms is the increased activity of STN before any significant striatal dopamine loss which leads to excess excitatory drive from STN to remaining SNc cells to restore the dopamine loss due to initial cell loss (Bezard et al., 1999; Vila et al., 2000; Obeso et al., 2004). This excess excitatory drive from STN eventually leads to excitotoxicity in SNc neurons. To overcome this excitotoxicity, subthalamotomy had to be applied very early after diagnosis of PD to have any neuroprotective effect (Guridi and Obeso, 2015; Guridi et al., 2016). The neuroprotective effect of dopamine restoration and subthalamotomy therapies are similar that is more effective in the early stages compared to the late stages of disease progression.

#### Deep brain stimulation

In our modeling study, we have explored various aspects of DBS protocol from stimulus waveforms to stimulus configurations and other optimal DBS parameters. From the simulation results, it can be suggested that biphasic stimulus waveform with four-contact point stimulation configuration showed maximal neuroprotective effect since biphasic stimulus guarantees charge-balance in the stimulated neuronal tissue (Hofmann et al., 2011) and DBS parameters were given in the *Table-2*.

It has been reported that long-term stimulation (DBS) of STN results in slowdown of the progression of SNc cell loss in animal models (Benazzouz et al., 2000; Maesawa et al., 2004; Temel et al., 2006; Wallace et al., 2007; Spieles-Engemann et al., 2010; Musacchio et al., 2017), but the mechanism behind the neuroprotective benefits of DBS is not clearly elucidated. In order to understand the neuroprotective effect of DBS in PD, we have investigated three prominent hypotheses namely excitation, inhibition and disruptive actions of DBS (McIntyre et al., 2004; Chiken and Nambu, 2015). In the excitation hypothesis, only DBS was applied which results in increased firing rate in STN and leads to more excitatory drive to SNc which eventually kills the SNc cells due to stress. Therefore, considering only the excitation hypothesis cannot explain the neuroprotective effect of DBS. Next, inhibition hypothesis was implemented where antidromic activation of GPe result in the increased inhibitory drive to STN (Mandali and Chakravarthy, 2016). In this scenario also, the neuroprotective effect of DBS could not be comprehensively explained. Finally, the disruptive hypothesis was implemented by increasing the axonal and synaptic failures in STN population during DBS therapy (Rosenbaum et al., 2014). From simulation results, it was observed that the progression of SNc cell loss kept on delaying as the percentage of STN axonal and synaptic failure increases. Therefore, it can be inferred that DBS blocks the propagation of pathological oscillations occurring in STN to other nuclei; in order words DBS disrupts the information transfer through the stimulation site, producing neuroprotection effect in SNc (Ledonne et al., 2012; Rosenbaum et al., 2014; Chiken and Nambu, 2015).

### A Major prediction from this modeling study

The proposed model was able to predict the influence of STN and sensitivity of stress threshold in causing excitotoxicity in SNc where STN activity and variability in stress threshold might be the cause of excitotoxicity. From the neuroprotective point of view, our model predicts that glutamate inhibition is more effective compared with dopamine restoration and subthalamotomy therapies in the late-stage of disease progression. From our study, we suggest that DBS protocol with four-contact point stimulation with biphasic stimulus waveform showed maximum neuroprotective benefits in SNc. We are also predicting the mechanism behind the neuroprotective effect of DBS where the information transfer through the stimulated site is blocked by synaptic depression in STN, which is consistent with others findings (Rosenbaum et al., 2014).

### Suggesting further experiments

By using the primary culture of SNc neurons (Gaven et al., 2014), the effect of glutamate on the SNc cells under normal and energy-deficient conditions can be studied (Novelli et al., 1988). Using brain slice cultures or co-culture systems that comprise of SNc-STN system, one can investigate the conditions under which excitotoxic runaway effect is observed (Renaud and Martinoli, 2016).

### Limitations and Future work

The timescales which are represented in the results of the proposed model are not realistic, as the neurodegeneration which occurs over the years in PD was exhibited in a few tens of seconds in the model. This limitation is inevitable in computer simulations since it is impractical to simulate for months and years. The difficulty arises due to the fact that the simulation must span widely separated time scales – sub-millisecond time scales to describing spiking activity and years to describe neurodegenerative processes.

The major inputs to the SNc neurons come from the striatum which is not included in the model. As, our objective was to investigate the extent of STN-mediated excitotoxicity in SNc, we avoided any other structures which can influence this phenomenon at present.

In the proposed model, the variability of stress threshold, which is analogous to an apoptotic threshold (that can be broadly associated with the available energy represented as 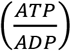 ratio (Wang and Michaelis, 2010)) is sensitive enough to alter the model results is a constant parameter. We would like to allow this parameter to vary from neuron to neuron in the SNc population. This can be achieved by introducing models of astrocyte and vascular networks that can influence the stress threshold. With the astrocyte layer introduced, the effect of astrocytes on the therapeutic effect of DBS can be explored (Fenoy et al., 2014).

In the future, we will incorporate a detailed biophysical model of SNc along with dopamine synthesis pathway (Tello-Bravo, 2012), apoptosis pathway (Hong et al., 2012) and neural energy supply-consumption properties (Wang et al., 2017) to the current model. The synaptic weights in the proposed model are not dynamic, we would like to include some type of learning principle by incorporating STDP type of learning rule in STN population which can show the long-term effect of DBS treatment (Ebert et al., 2014; Iakymchuk et al., 2015). The bigger goal is to incorporate detailed SNc module into a large-scale model of basal ganglia (Muralidharan et al., 2016) in understanding the effect of therapeutics on the behavioral response (Érdi and Tóth, 2005; Aradi and Érdi, 2006; Erdi et al., 2006; Kiss and Érdi, 2006).

Our hypothesis behind this whole study is to understand the pathogenesis of PD as cellular energy deficiency in SNc as a cause. As Wellstead and Cloutier pointed out (Wellstead and Cloutier, 2011), PD should be understood by placing brain energy metabolism as a core module and other cellular processes related to PD can be incorporated thereby studying it in an integrative environment (see the fig. 12 in (Wellstead, 2010)).

